# Induction of hemodynamic traveling waves by glial-related vasomotion in a rat model of neuroinflammation: implications for functional neuroimaging

**DOI:** 10.1101/2024.12.13.628348

**Authors:** Mickaël Pereira, Marine Droguerre, Marco Valdebenito, Louis Vidal, Guillaume Marcy, Sarah Benkeder, Paul Marchal, Jean-Christophe Comte, Olivier Pascual, Luc Zimmer, Benjamin Vidal

## Abstract

**Background:** Cerebral hemodynamics are crucial for brain homeostasis and serve as a key proxy for brain activity. Although this process involves coordinated interaction between vessels, neurons and glial cells, its dysregulation in neuroinflammation is not well understood.

**Methods:** We used in vivo mesoscopic functional ultrasound imaging to monitor cerebral blood volume changes during neuroinflammation in male rats injected with lipopolysaccharide (LPS) in the visual cortex, under resting-state or visual stimulation, combined to advanced ex vivo techniques for glial cell reactivity analysis.

**Findings:** Cortical neuroinflammation induced large oscillatory hemodynamic traveling waves in the frequency band of vasomotion (∼0.1 Hz) in both anesthetized and awake rats. Vasomotor waves traveled through large distances between adjacent penetrating vessels, spanning the entire cortex thickness, and even extending to subcortical areas. Moreover, vasomotion amplitude correlated with microglial morphology changes and was significantly reduced by astrocytic toxins, suggesting that both microglia and astrocytes are involved in the enhancement of vasomotion during neuroinflammation. Notably, functional connectivity was increased under this oscillatory state and functional hyperemia was exacerbated.

**Interpretation:** These findings reveal new spatiotemporal properties of cerebral vasomotion and suggest this is a major component of brain hemodynamics in pathological states. Moreover, reactive microglia and astrocytes are participating to increased vasomotion during neuroinflammation. For the field of functional neuroimaging, our results advocate for considering 0.1 Hz hemodynamic oscillations as an important complement to traditional measurements, particularly in neuroinflammatory conditions. Indeed, brain hemodynamics may provide insights not only into neuronal activity but also glial reactivity.

**Funding:** Supported by ANR (“LabCom-NI2D”) and Auvergne-Rhône-Alpes Region (“BI2D”).

## Introduction

Cerebral hemodynamics constitute a fundamental component of the central nervous system (CNS) by providing the necessary oxygen and nutrients to support its substantial energy consumption. Cerebral blood flow is dynamically adapted in response to changes in neuronal activity, a process known as neurovascular coupling. This fundamental process underlies functional neuroimaging techniques that measure hemodynamic parameters to infer changes in brain activity, such as functional magnetic resonance imaging (fMRI), intrinsic optical imaging, laser speckle contrast imaging, near-infrared spectroscopy, and more recently functional ultrasound imaging (fUSi)^1–6^. It is therefore of major importance to understand the processes governing cerebral hemodynamics to gain deeper insights into brain functions, as well as to accurately interpret functional neuroimaging findings, in both physiological and pathological states.

Despite many efforts and advances, the cellular mechanisms that govern hemodynamic fluctuations in resting-state as well as functional hyperemia (i.e, the increase of cerebral blood flow in response to an increased neuronal activity) remain elusive and controversial. An emerging hypothesis is that vasomotion, i.e., the rhythmic fluctuations of vessel diameter (∼0.1 Hz) produced by the autonomous contraction of smooth muscle cells, is naturally synchronized to neuronal activity to meet the energetic demands under physiological conditions^7^. This model, although debated^8^, provides a possible cellular basis for functional connectivity. In parallel, other studies have suggested that glial cells are equally important for regulating hemodynamic fluctuations. Astrocytes, in particular, are strategically positioned to mediate interactions between neurons and blood vessels as their end-feet envelop most of the cerebral vasculature. Astrocytic control of cerebral blood flow during functional hyperemia has been extensively studied, and involves the Ca^2+^-dependent release of vasoactive metabolites of arachidonic acid, as well as potassium efflux from the endfeet^9–16^. They also appear to be involved in the regulation of cerebral hemodynamics during resting-state, as they were recently suggested to exert a negative control on the amplitude of vasomotion at 0.1 Hz in arterioles^17^. This brings vasomotion at the center of a hypothetic cellular model for explaining spontaneous hemodynamic fluctuations and resting-state connectivity, with SMCs setting the tempo around 0.1 Hz, astrocytes calibrating the gain, and neurons serving as the conductor of this orchestrated response. Other glial actors such as microglia, the key regulators of inflammatory processes in the brain, may also be involved as they can dynamically interact with the neurovascular unit and have been shown to modulate functional hyperemia^18^.

Fewer studies have focused on the cellular mechanisms involved in the control of cerebral blood flow in pathological conditions^9,19^. This is important because all cellular players of the neurovascular unit, and in particular microglia and astrocytes, undergo major morphological, functional and molecular changes during inflammatory processes that can occur in many neuropsychiatric conditions. Such changes could have major impact on the regulation of the cerebral hemodynamics, which must be taken into account for the interpretation of neuroimaging studies and for understanding the alterations of brain homeostasis that can occur during neuroinflammatory processes .

Using functional ultrasound imaging (fUSi), we aimed to extensively characterize the cerebral hemodynamics changes that occur in a neuroinflammatory state and the influence of reactive glial cells in these alterations. This recent technique can map cerebral blood volume (CBV) changes at a mesoscopic scale with higher sensitivity than fMRI^20^ even in deep areas, unlike optical techniques, and can be used in freely moving rodents^21–23^. CBV changes were studied during LPS-induced neuroinflammation through injection in the primary visual cortex, during resting-state as well as visual stimulation. In parallel, the impact of the inflammatory challenge on the reactivity of glial cells was assessed by advanced morphological analyzes and single-nuclei transcriptomics. We show that cortical neuroinflammation can drastically enhance hemodynamic oscillations at 0.1 Hz, the typical rhythm of vasomotion. Adding to the recent descriptions of vasomotion as long-wavelength traveling waves along cortical pial vessels^24^, the vasomotor waves traveled across large areas between neighboring penetrating vessels and even reached subcortical vessels. We also highlight that vasomotion amplitude may be locally driven by the presence of reactive glial cells, as suggested by a correlation between microglial morphological changes and hemodynamic oscillations at 0.1 Hz, and the significant inhibition of inflammatory vasomotion by astrocytic toxins. Finally, enhanced vasomotion was associated with an increase in cortical connectivity, extending previous findings that vasomotor fluctuations are a major component of functional connectivity, and with an increased hyperemic response during visual stimulation, suggesting that the two phenomena share common but non-competitive pathways under specific states. These findings demonstrate that vasomotion and traveling waves around 0.1 Hz are fundamental players of cerebral hemodynamic fluctuations, especially during neuroinflammation, and emphasize the importance of reactive glial cells in the control of spontaneous changes of cerebral blood volume in pathological states, usually interpreted only in terms of neuronal activity in functional neuroimaging studies.

## Methods

### Ethics

All animal experimentation and housing were carried out in accordance with the guidelines of the European Union (Directive 2010/63/EU) and approved by the French Ministry of Research and CELYNE ethical committee (authorization reference: APAFIS-36043). All efforts were made to ensure the animal welfare.

### Animals

The animals were kept under controlled environmental conditions (12/12 h light-dark cycle, food and water ab libitum, controlled temperature) with at least one week of acclimatation before any experiment. A total of 62 Wistar male rats (from Janvier Labs, Le Genest-St-Isle, France) were used in this study, weighing between 225-250g (approximately 6 weeks of age) at arrival to the laboratory. Females were not used due to possible interactions between estrous cycle and inflammation, which restricts the relevance of the study to male rats. The animal well-being was monitored every day following surgery using a score sheet. Humane endpoints were defined in terms of weight loss, physical appearance (grooming, orbital tightening, posture), and general behavior. These criteria would have led to euthanasia if any of the signs had become severely impaired (no such cases were observed during the study, but occasional weight loss or rare signs of discomfort were addressed respectively with supply of food softened into a mash and administration of buprenorphine).

For the main experiment, 40 rats were used for a protocol initially designed to explore the consequences of inflammation and astrocytic lesions on brain hemodynamics (protocol was not preregistered). An initial sample size of 6 was defined based on effect size and variability observed in previous studies from our laboratory. Animals were allocated to experimental groups in a balanced manner across batch numbers, week of experiments and groups; no formal randomization tool was used. The rats first underwent a skull thinning surgery the day before beginning the imaging protocol (6 animals were excluded after the surgery due to the presence of hematomas). The day of the first fUSi session, the rats received intracerebral bilateral injections of the following agents: NaCl 0.9% (VEH, n = 6), lipopolysaccharide (LPS, n = 6), L-alpha amino acid (AAA, n = 6), fluorocitrate (FC, n = 5), a combination of LPS and AAA (LPS+AAA, n = 5), or a combination of LPS and FC (LPS+FC, n = 5), 1 h before the fUSi acquisition under anesthesia (except one rat in the LPS+AAA group that died before starting the imaging session). One resting-state from AAA-1h group was excluded due to inconsistency of acquisition parameters compared to the rest of the study. The rats were then used for a second fUSi session under anesthesia, 48 h after the intracerebral injections. The brains of all rats imaged twice were used for immunofluorescence staining (n = 6 for VEH, LPS and AAA; n = 5 for FC, LPS+AAA and LPS+FC) for assessment of different neuropathological features and verification of astrocyte loss in the groups that received the gliotoxins (leading to the exclusion of one rat from the LPS+AAA group because no astrocytic lesion could be detected for this animal). Some rats were not used for the immunofluorescence analyses due to staining or tissue low quality in the areas of interest.

Additional studies were planned to enhance the findings interpretability. For further evaluation of the temporal effects of LPS, 12 additional rats underwent thinned-skull surgery (24 h before imaging and/or intracerebral injection) and were used for one anesthetized fUSi session, either without injection (n = 3), 24 h (n = 3) or 7 days (n = 3) after LPS injection. For awake fUSi, 4 other rats underwent skull thinning (24 h before imaging and/or intracerebral injection) and were imaged in freely moving protocol before LPS injection (n = 3), 1 h (n = 1) and/or 48 h (n = 3) after bilateral LPS injections. For single-nuclei transcriptomic analysis, 6 other rats were bilaterally injected with VEH or LPS and euthanized before sampling the brains 48 h (n = 2 VEH, n = 2 LPS) or 7 days (n = 2, LPS) after injection.

### Surgical procedures

#### Skull Thinning

A thinned-skull window was performed before fUSi to enable sufficient penetration of acoustic waves, as previously described in the rat^25,26^. For this purpose, rats were anesthetized with isoflurane (induction 4%, then 2% in a 50/50 mix of air and O_2_ during the surgery) and received a subcutaneous injection of buprenorphine (0.05 mg/kg; Buprecare, Axience). The top of the head was shaved and cleaned with povidone-iodine (Vetedine, Vetoquinol), then 30 min after buprenorphine pretreatment, the scalp was incised. The skin was carefully pulled to visualize the lateral sides of the skull from bregma to lambda anatomical landmarks. The skull thinning was performed from -3.00 mm to -8.00 mm (AP) and +5.00 to 5.00 mm (L) from bregma using a drill at low speed with a micro drill steel burr (Harvard Apparatus, 75-1887). To avoid overheating during drilling sessions, cold saline solution was regularly added on the skull. When the skull was thin enough to be flexible, the scalp was sutured, povidone-iodine and lidocaine 5% (EMLA 5%, Aspen) were locally applied, and the rats were placed under heating light to recover from anesthesia.

#### Intracerebral injections

After 24 h of recovery, rats were anesthetized with isoflurane (4% for induction, then lowered to 2% in a 50/50 mix of air and O_2_ during the surgery) and placed in a stereotaxic frame with a heating blanket. The wound was cleaned and lidocaine 2.5% was applied before removing sutures and exposing the thinned-skull window. Animals received bilateral injections of 4 µl of solution (LPS, FC, AAA, VEH, LPS+AAA or LPS+FC) at a rate of 0.4 µl/min into the primary visual cortex (AP: -6; L: ±4.0; D: 1.6 - in mm from bregma). At the end of each injection, the needle was left in place for 5 min before being slowly removed. LPS (1.25 µg/µl; L2880), FC (0.25 nmol/µl; F9634) and AAA (12.5 µg/µl; A7275) were purchased from Sigma-Aldrich. For the preparation of FC solution, DL-fluorocitric acid barium salt was dissolved in 1 mL of HCl 0.1 M. Two drops of Na_2_SO_4_ were then added to precipitate Ba^2+^. After adding 2 mL of Na_2_HPO_4_, the solution was centrifuged at 1000 g for 5 min, and 0.31 mL of supernatant was diluted in 0.9% NaCl to the final concentration (while adjusting the pH to 7.4).

### Functional Ultrasound Imaging

Rats were imaged using a preclinical ultrasound imaging system (Iconeus V1, Paris, France). Doppler vascular images were obtained using the Ultrafast Compound Doppler Imaging technique^27^. Each frame corresponded to a Compound Plane Wave frame^28^ resulting from the coherent summation of backscattered echoes obtained after successive tilted plane waves emissions. Each stack of 200 compounded frames acquired at 500 Hz frame rate was processed with a dedicated spatiotemporal filter using Singular Value Decomposition^29^ to extract the blood volume signal from the tissue signal, to obtain the Power Doppler images. These final images have a temporal resolution of 0.4 s, a spatial resolution of 100 µm and are directly proportional to the cerebral blood volume.

#### Anesthetized imaging protocol

At the end of the intracerebral injection procedure, the animals received a subcutaneous injection of medetomidine (0.05 mg/kg; Domitor, Vetoquinol), and were moved in another stereotaxic frame under the ultrasound imaging probe before performing the acute fUSi acquisition, with isoflurane being still delivered at 2% through a mask to maintain anesthesia. A subcutaneous catheter was placed and the medetomidine infusion (0.1 mg/kg/h) was started, while progressively decreasing isoflurane to 0.4%, to follow the consensus protocol for studying functional connectivity in the rat^30^. Respiratory rate and body temperature were continuously monitored (TCMT, Minerve, France) and the animal temperature was maintained using a heating blanket. The thinned skull was exposed and covered with ultrasound gel. After reaching the final concentration of isoflurane, a period of at least 10 min was added before starting the imaging experiments to ensure physiological stability of the animals. Then the acute session of fUSi was performed, that included a resting-state acquisition followed by a visual stimulation acquisition. At the end of this imaging session, the scalp was sutured, and betadine and lidocaine were applied on the wound. Finally, rats received a subcutaneous injection of atipamezole (5 mg/kg; Antisedan, Vetoquinol) and were placed under heating light to recover from anesthesia. 48 h later, the second, delayed fUSi session was performed. Animals were anesthetized first with isoflurane under the same conditions and duration as for the stereotaxic injections performed 2 days before, and then were placed in a stereotaxic frame under the ultrasound imaging probe for the second session under the same anesthesia protocol (combination of isoflurane 0.4% and subcutaneous medetomidine, bolus 0.05 mg/kg followed by infusion at 0.1 mg/kg/h), that also included a resting-state acquisition followed by a visual stimulation. At the end of this second session, an ultrasound localization microscopy (ULM) acquisition was also performed.

#### Resting-state protocol

For resting-state fUSi, the probe was placed above the injection site (AP: -6; L: ±4.0; D: 1.6 - in mm from bregma) to follow the CBV changes in the primary visual cortex during a 20-minute period, with a sampling rate of 0.4 s.

#### Visual stimulation protocol

For fUSi acquisition with visual stimuli, the probe was kept in the same coronal plane. The visual stimuli protocol was performed using a screen placed in front of the animal, as previously described^31^. Briefly, the acquisition began with an initial 30-second rest period in the dark, followed by 4 blocks of stimuli, all separated by a 45-second rest period in the dark. Each visual stimuli block consisted of a 30-second period with black and white flickering at 3 Hz, as this frequency was shown to elicit the best response in the visual cortex in a previous fUSi study in the rat^32^. The stimulation was performed using a homemade Python script.

#### Ultrasound localization microscopy protocol

At the end of the last imaging session, an ULM acquisition was performed for each animal. An intravenous catheter was set into the caudal vein before starting the acquisition. Two boluses of 300 µL of a microbubble ultrasound contrast agent (Sonovue, Bracco Imaging, France) were slowly injected at the start of acquisition and at 2.5 min of acquisition, for a total acquisition time of 5 min. This sequence enables to visualize the vasculature and to measure the blood flow velocity with very high spatial resolution (10 µm)^33^.

#### Kinetics of LPS effect

To further evaluate the temporal effects of LPS, an additional group of animals underwent a resting-state acquisition after 0, 1 or 7 days following the intracerebral injection of LPS using the same imaging, anesthesia and surgery conditions as stated above. The impact of skull thinning on our results was also assessed with three rats without injection of LPS, scanned in fUSi under the same conditions.

#### Freely moving awake imaging protocol

A resting-state fUSi protocol was also performed in awake freely-moving rats. These experiments included a first phase of handling/habituation of 5 days, followed by a surgery to implant the head plate (designed by Iconeus, Paris, France). After a minimal period of 24 h of recovery, a 3-days habituation period was performed, during which the rats were free to move between their home cage and the cage used for imaging experiments. 24 h later, a first baseline fUSi session was performed. Then, the bilateral stereotaxic injections of LPS or VEH were performed as stated previously in primary visual cortex (AP: -6; L: ±4.0; D: 1.6 - in mm from bregma), followed by the second resting-state fUSi in awake condition, 1 h after the intracerebral injections. Animals were left in their initial cage for 2 days before their last resting-state imaging session.

### Single-nucleus transcriptomics

#### Single-nuclei isolation

Dissected frozen cortex was resuspended into 600µl of cold homogenization buffer that consisted of 10mM Tris Hcl pH 7.5, 146mM NaCl, 1mM CaCl_2_, 21mM MgCl_2_, 0.03% Tween20, 0.01% BSA, 10% EZ buffer (Sigma) and 0.2U/µl Protector RNase Inhibitor (Roche). Tissues were then transferred into 2mL dounce (Kimble) and homogenized using 10 strokes of the loose pestle followed by 8 strokes of the tight pestle to release nuclei, on ice. Homogenate was then strained through a 70μm cell strainer (Pluriselect) and centrifuged at 500g for 5 minutes to pellet nuclei. After removing supernatant, nuclei were washed in 1mL resuspension buffer containing 10mM Tris Hcl pH 7.5, 10mM NaCl, 3mM MgCl_2_, 1% BSA and 0.2U/µl Protector RNase Inhibitor and centrifuged at 500g for 5 minutes. Nuclei were then fixed with the Evercode Nuclei Fixation kit (#ECF2003) according to manufacturer instructions (Parse Biosciences) and FACS-sorted based on DAPI intensity to remove debris (BD ARIA). Nuclei were then frozen down.

#### Single-nucleus RNA sequencing

Nuclei were thawed and loaded in round 1, 2 and 3 plates for combinatorial barcoding according to manufacturer instructions (#CW02030, Parse Biosciences). Six sub-libraries of 12,500 nuclei were prepared to target 75,000 nuclei and 12,000 nuclei per condition. Libraries were sequenced using the Novaseq 6000 platform (Illumina) to target 50,000 reads per nucleus. Parse demultiplexing pipeline v1.1.2 was used to align reads on the rat reference genome (Rattus_norvegicus.mRatBN7.2) and to produce the count matrix.

### Immunofluorescence

Rats were deeply anesthetized and transcardially perfused with 4% paraformaldehyde in phosphate-buffered saline (PBS). Brain tissues were post-fixed in 4% paraformaldehyde for 24 h, then immersed in 20% sucrose for 48 h. Fixed brain sections were cut at 40 µm using a cryostat (Leica Biosystems – CM3050S). After washing in PBS, the sections were permeabilized in PBST-BSA (0.5% Triton X + 5% BSA in PBS) for 90 min. Primary antibodies, diluted in blocking solution, were applied overnight at 4°C. After three washing in PBS, appropriate secondary antibodies coupled to fluorochromes were diluted in PBST-BSA and applied for 1 h at RT. Slides were rinsed and coverslipped with DAPI fluoromount mounting media. The antibodies list and RRID tags is provided in the Supplementary Table 1.

### Data analysis

*fUSi.* All data processing steps were performed as previously described^34,35^. Briefly, a custom template was created by a first automatic registration step and by computing the mean of all fUSi acquisitions gathered during resting-state and visual stimulation. This template enabled to manually refine the registration of all images to each other.

#### Resting-state

For the analysis of CBV changes and static functional connectivity, the registered images were band-pass filtered between 0.0008 Hz and 0.2000 Hz. Then, a 2D slice corresponding to the imaging plane (−6 mm from bregma) was extracted from the SIGMA atlas^36^ and manually co-registered on the fUSi template in the ITK-Snap software. The Power Doppler signal was automatically extracted from each region of interest - namely, the primary and secondary visual cortices (PrimVisCtx and SecVisCtx), the retrosplenial dysgranular cortex (RetroDysCtx), the parts A and B of the retrosplenial granular cortex (RetroGrCtx_A_ and RetroGrCtx_B_) and the superficial part of superior colliculus (SuperiorColl) for each individual scan using a Matlab script. For the functional connectivity analysis, the temporal correlations were computed using the Pearson correlation coefficient for each pair of regions. Fisher Z transformation was applied to the correlation coefficients before calculating the differences between the correlation matrices in each condition relative to the vehicle.

For the analysis of the amplitude of specific frequency bands (also known as fALFF for fractional amplitude of low-frequency fluctuations), no frequency filtering was done before applying the slice of atlas on the registered images for extraction of the fALFF in the different ROIs. fALFF was obtained using a custom Matlab script adapted from the REST toolbox^37^. Briefly, the power over a frequency band of interest (0.08 to 0.15 Hz) was calculated for each pixel and divided by the power of the entire frequency range.

For the principal component analysis, the images were band-pass filtered between 0.0008 Hz and 0.2000 Hz, centered by subtracting the global mean, and normalized prior to reshaping each acquisition into two dimensions. Following the approach of Cabral et al.,, we calculated and then averaged the covariance matrices of all acquisitions for each group to capture common variance patterns^38^. The eigenvectors were derived from this mean covariance matrix and sorted in descending order of their corresponding eigenvalues. The first principal components’ spatial maps were visualized by reshaping each eigenvector into the original image format (only the components with eigenvalues higher than 0.1 were retained, with up to 10 components for simplification). The temporal patterns associated with these components were obtained by projecting the time series data onto the eigenvectors. The Fourier transform was applied to these temporal patterns to calculate their amplitude spectra, which were then averaged across acquisitions. For visualization, some acquisitions were reconstructed using the first 10 principal components, summing the product of each principal component and its corresponding temporal pattern. The scree plot, showing the proportion of total variance explained by each principal component, was achieved by computing the total variance across all components using the global sum of eigenvalues and calculating the cumulative sum of the individual variances explained by the components as a proportion of the total variance, in descending order of explained variance.

For the awake dataset, the same preprocessing and analysis (functional connectivity matrices, fALFF and PCA) were performed on selected continuous periods of 120s that were devoid of movement artifacts for each scan (estimated using a threshold defined on the time-courses of the Power Doppler signal, and further confirmed by the absence of obvious artifacts in the spatial maps of the principal components).

For the analysis of traveling waves, the images were filtered between 0.08 Hz and 0.15 Hz to focus on the frequency of oscillations. We used the AquA software^39^ to automatically extract static and traveling events. This software detects signal peaks propagating across space without predefined assumptions about event locations or timings. Originally developed for fluorescence imaging and astrocytic calcium waves, AquA is explicitly designed for working with various biological contexts and spatiotemporal scales. After normalization to correct for the baseline differences in signal intensity and estimate the noise variance, the groups of active pixels were found using an intensity threshold of 20, a smoothing parameter of 0.5 and a minimum size criterion of 15 pixels. Significant pixels (z-score above 5) were retained, and events were detected by identifying local maxima (seeds) in these pixels, extending them in space and time to estimate their propagation. Events were further refined using a z-score threshold of 5, based on peak size and noise level. One acquisition from VEH-1h group was discarded due to failure of AquA to complete its analysis. We classified events as dynamic/traveling if their propagation distance (defined by the software as the sum of the changes of centroid locations from the beginning to the end of event) exceeded 5 pixels, and as static otherwise. We averaged the different features of interest across all dynamic events for each scan for group comparisons (percentage of dynamic events relative to the total number of events, and specifically for traveling events, amplitude of peaks and propagation corresponding to the distance crossed from the beginning to the end of the event). To explore traveling wave trajectories and locations in the different groups, we clustered the propagation maps generated by AquA (total of 17377 events with an area > 5 pixels). The individual maps were first binarized, reshaped and concatenated in a two-dimensional array, followed by singular value decomposition (SVD) to retain the first 3000 components. We then performed k-means clustering to identify 4 main clusters corresponding to different spatial locations, regardless of the direction of the waves. These clusters were further subclustered based on horizontal and vertical propagation directions, obtained by averaging the delay values of the original propagation maps in these directions and concatenation of the directional vectors into a two-dimensional array, followed by k-means clustering. This process allowed us to identify 6 distinct spatiotemporal patterns in our dataset, in addition to the events that were static or followed an undefined propagation pattern.

The dynamic seed-based connectivity results were computed on 0.08-0.15 Hz band-pass filtered representative individual scans, with a window size of 15 successive images and a step of 1 image, by measuring the Pearson correlation coefficients between the average time course in the right visual cortex as a seed and all pixels of the image. The time course of the average correlation coefficients in the contralateral hemisphere was displayed as a heatmap with superimposed average CBV changes occurring in both hemispheres.

#### Visual stimulation

For visual stimulation analysis, the average CBV changes during all stimulations were extracted in the regions of interest and the four time-courses corresponding to each stimulation block were averaged for all individuals. The mean CBV changes values during stimulation as compared to the baseline were estimated for each individual before group comparisons.

#### Ultrasound localization microscopy

The speed maps were reconstructed using a dedicated Iconeus code, based on the tracking of individuals microbubbles in the ULM data. An additional spatial filter was applied to remove outliers, which decreased the spatial resolution to 10 × 10 x 400 μm. The analysis was restricted to the primary visual cortex area. The pial vessels were manually drawn on each individual image on the absolute speed map, and the penetrating vessels were further differentiated as arterioles or venules depending on the vertical direction of the flow indicated in the directional map. The absolute speed values were averaged in these different vascular compartments for each individual scan and compared between groups.

#### Immunohistochemistry

Slides were scanned with ZEISS Axioscan 7 Microscope Slide Scanner at the East Lyon Quantitative imaging center of University Lyon1 (CIQLE). For quantification of Iba1, GFAP, and NDRG2 markers, the mean intensity was measured via ImageJ in four to eight areas in the primary visual cortex for each animal (after normalizing the intensity to that of the whole image for NDRG2). For NeuN counting, the neuron number was automatically determined using Spot detector tool of the Icy software.

#### Microglia morphological analysis

For the comparison of microglial morphology in the injection site between all groups, a total of 250 to 400 microglia were cropped individually randomly in four to eight areas in the primary visual cortex for each group (excluding 2 animals that did not display sufficient signal quality). Between 400 and 560 microglia were additionally cropped in two other subcortical areas from the LPS and VEH groups, given the presence or absence of oscillations

The first subcortical zone consisted of regions from the hippocampus and dorsal midbrain showing oscillations. The second subcortical zone represented regions devoid of oscillations from the ventral midbrain. The crops were then binarized and their morphology automatically analyzed with a recently developed pipeline called MorphoCellSorter^40^, using Matlab R2018. Briefly, 20 non dimensional morphological indexes are measured. Then, a principal component analysis (PCA) is performed to filter the parameters and select only the most discriminant ones. The selected parameters and their weight on the first principal component are subsequently used to form a derivative of an Andrews plot, which allow the two-dimensional representation of high dimensional data. In this representation, each curve represents one microglia, and cells having similar morphological characteristics follow similar trajectories. It is used to establish a ranking of the cells according to their morphological features by using the Andrews score, corresponding to the value of the Andrews plots at the time point where the curves are the most dispersed (i.e. when the variance is maximal).

#### Single-nucleus RNA-seq data analysis

The gene expression matrix from Parse pipeline was used for downstream analysis using the software R (version 4.4.0) and the R toolkit Seurat (version 5.1.0). Nuclei were excluded from downstream analysis when they had more than 1% mitochondrial genes, fewer than 100 unique genes, more than 15,000 unique molecular identifiers (UMIs) and detected as doublets using AUC cell type marker scores based on AUCell R package. A total of 64,471 nuclei were selected for the three groups and 6 animals studied. Gene expression was normalized using the standard Seurat workflow and the 2000 most variable genes were identified and used for principal component analysis (PCA) at 50 dimensions. The top most significant principal components (PCs) were selected for generating the UMAP, based on the ElbowPlot method in Seurat. Clustering of cells was obtained following Seurat graph-based clustering approach with the default Louvain algorithm for community detection. We then performed differential expression analysis using the FindMarkers function of Seurat and annotated clusters based on expression of marker genes. We thus manually annotated the major classes of cells: Neurons, Microglia, Astrocytes, Oligodendrocytes (OLs) and Vascular cells.

#### Gene set enrichment

Overrepresentation analyses of biological processes (BPs) were performed using the R package “Cluster Profiler”^41^. DEGs were extracted using the following criteria: Logfc.threshold = 0.25, min.pct = 0.1.

### Statistical analysis

All statistical analyses were performed using GraphPad Prism 8.0.1, except the GLM analyses that were performed using R. For all experiments, a level of p < 0.05 was accepted as evidence for a statistically significant effect (*p < 0.05; **p < 0.01; ***p < 0.001). As a general rule, unpaired t-tests, one-way or two-way ANOVAs were chosen for all analyses if data followed a normal distribution and showed homogeneity of variance, verified using respectively Shapiro-Wilk and Levene tests (p>0.05 for all analyses, Bonferroni corrected). If the required assumptions were not met, non-parametric tests were applied in Graphpad Prism (or GLM in R for alternatives to two-way ANOVAs with more than two experimental groups). For the functional connectivity analyses, the different groups were compared using two-way ANOVAs followed by Sidak’s correction for multiple comparisons. For the fALFF analyses, Mann-Whitney tests with false discovery rate (FDR) correction of 5% were used for comparing VEH and LPS, and a GLM analysis (glm function in R, fALFF ∼ Group*Region, gamma family) with Dunnett’s correction was used for other analyses involving more than two groups. For the comparisons of the ROIs time-courses during stimulation, two-way repeated-measures ANOVAs were used followed by Sidak’s multiple comparisons tests (p < 0.05). For ULM analysis, the difference of perfusion was assessed using Mann-Whitney tests with FDR correction of 5%. For the traveling waves analysis, the general features of dynamic events were calculated in AquA software, averaged for each individual, and compared using one-way ANOVA followed by Dunnett’s test or unpaired t-test depending on the number of groups. For comparison of the occurrence of different wave patterns among groups, the percentages of the different patterns were compared between each group and the vehicle using two-way ANOVAs for each time-point, and between each time-point for each group using mixed-effect analysis. The p-value of the interaction between the condition factor (group or time-point) and the patterns factor was used to identify the significant changes of propagation patterns. For histological analyses, expression of GFAP, NDRG2, NeuN and Iba1 were compared between groups using Kruskal-Wallis tests with FDR correction of 5%.

## Results

### Inflammation induced local hemodynamic oscillations at 0.1 Hz and increased functional connectivity

The effects of bilateral intracortical injection of LPS in the visual cortex were monitored in a coronal plane centered around the injected area using fUSi, first during resting-state at different times post-injection (Fig. 1a). The spontaneous CBV changes were extracted from different ROIs (Fig. 1b). No obvious modification of CBV changes was observed 1h or 48h after vehicle injection (exemplified in Fig. 1c). However, the administration of LPS induced a high-amplitude low frequency oscillatory pattern that appeared 48-h post-injection (Fig. 1d; sky-blue curves, as compared to the dark-blue curves at 1h). From the frequency spectrum, these waves are occurring between 0.08 and 0.15 Hz, on average centered around 0.125 Hz (Fig. 1e-f), the typical frequency band of vasomotion. To quantify this phenomenon, the frequency amplitudes between 0.08 and 0.15 Hz were computed for each individual scan and each group in the different ROIs. Whereas no significant difference were found between groups at 1h (all q > 0.9999 and mean rank differences of Mann-Whitney tests ranging from - 0.6667 to 1; Fig. 1g), the amplitude of oscillations between 0.08-0.15 Hz was significantly higher in LPS compared to vehicle at 48h post-injection (Fig. 1h) in the primary visual cortex (Mean rank diff = +4.667, q = 0.03636), and even more in the secondary visual cortex and the retrosplenial dysgranular cortex (Mean rank diff = +6.0000, q = 0.004545, for both). Indeed, when looking at the amplitude maps, oscillations occurred not only near the injection site but also spanned neighboring cortical areas, as shown in Fig. 1i. Functional connectivity variations between LPS and vehicle were also assessed for each time point (Fig. 1j). No difference was found during the first imaging session, 1h after injection (left diagonal). However, 48 h later (right diagonal), functional connectivity after LPS injection was significantly different (F (1,150) = 27.51, p < 0.0001, Two-way ANOVA, Sidak correction), with an increase between the secondary visual and the retrosplenial dysgranular cortices (Δr = +0.1654, p = 0.0374). Hence, the increase of functional connectivity was observed between highly oscillating cortical areas and neighboring areas.

**Figure 1:**
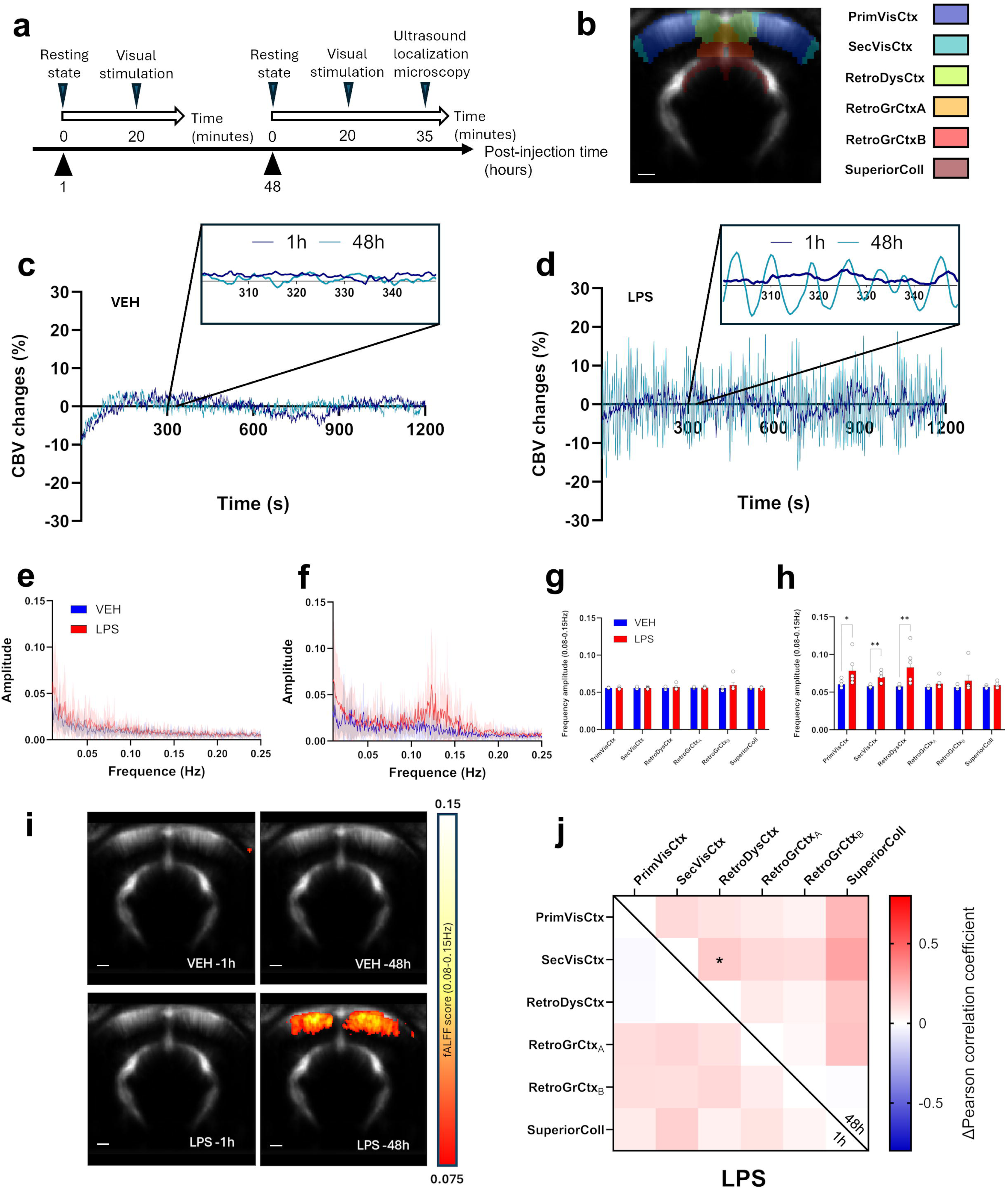
fUSi reveals spontaneous hemodynamic oscillations in anesthetized rats following the intracerebral injection of LPS. (a) Experimental timeline following the intracortical injection procedure. Each imaging session (1h and 48h post-injection) included a first resting-state acquisition followed by a second acquisition under visual stimulation. The second session was followed by an Ultrasound Localization Microscopy acquisition and animals were euthanized immediately after for brain sampling. (b) Mean of all registered fUSi images with overlay of the selected anatomical regions of interest from the SIGMA atlas. (c) Representative time curves of spontaneous CBV changes in the primary visual cortex of rats, 1h (dark blue) or 48h (light blue) following saline injection. (d) Same as (c) following LPS injection. The mean amplitude spectra (± SEM) after injection of LPS (red) or vehicle (blue) are shown for 1h (e) and 48h (f) post-injection. (g) Frequency amplitude in 0.08-0.15 Hz range at 1 h post-injection. Multiple Mann-Whitney tests followed by FDR correction of 5%, n=6 for both groups. (h) Frequency amplitude in 0.08-0.15 Hz range at 48 h post-injection. Multiple Mann-Whitney tests followed by FDR correction of 5%, n=6 for both groups. Each bar is the mean ± SEM. (i) Mean spatial maps of frequency amplitude in 0.08-0.15 Hz range in the different groups, with a threshold of fALFF>0.075. (j) Functional connectivity changes after 1 and 48 hours. Matrices of the correlation coefficient variations of LPS compared to vehicle are shown (bottom-left for 1 h post-injection, top-right for 48 h post-injection). For statistical comparisons, a Fisher Z-transformation was applied to the correlation coefficients before two-way ANOVA followed by Sidak’s multiple comparisons test (*p < 0.05; **p < 0.01; ***p < 0.001, ****p < 0.001). n=6 for both time points.

### LPS-induced hemodynamic waves are traveling through the cortex and subcortical regions

Next, we examined the dynamic changes occurring during inflammatory hemodynamic oscillations at 48 h after LPS injection. When measuring the dynamic functional connectivity between the left and right primary visual cortices, we observed an alternation between periods of hyper- and hypo connectivity towards extreme values (Fig. 2a) that were not observed at 1 h post-injection (Fig. 2b). A closer look at the individual maps of relative changes of CBV revealed that this was associated with the presence of hemodynamic traveling waves with variable spatial trajectories (See also Supplementary Movie 1). For illustration of this phenomenon, three representative time periods from Fig. 2a are displayed in terms of relative CBV maps (Fig. 2c), reflecting high correlation (time-period 1), no correlation (time-period 2) or anticorrelation (time-period 3) in one typical animal. Period 1 was associated with a fast bilateral cortical wave with successive decrease and increase of CBV that also involved subcortical regions. Period 2 involved unilateral waves with a high amplitude event that started from the left cortex to propagate to the left dorsal and then ventral hippocampus, while period 3 ended with a wave that traveled from the left hippocampus to the right retrosplenial and then right visual cortex, leading to the anticorrelation. We then used a peak detection software dedicated to the analysis of traveling events in stacks of 2D images to automatically identify CBV increases in our data and understand the trajectories of traveling waves (see Methods). The delay maps of all detected events (that is, local peaks of CBV) were classified in terms of spatial location and horizontal or vertical directionality by k-means clustering, enabling to reveal six different patterns of propagation (Fig. 2d). The traveling waves mainly involved adjacent penetrating vessels in the cortex, either from the retrosplenial cortex to the visual cortex (medio-temporal) or in the reverse direction (temporo-medial), in one hemisphere or bilaterally, or sometimes from one hemisphere to another (trans-hemispheric). In some cases, waves traveled from the cortex to the hippocampus (cortico-hippocampal) or reversely (hippocampo-cortical), or from the cortex to other subcortical regions (cortico-subcortical). This analysis also enabled to globally quantify the amount of traveling waves and their properties. Interestingly, dynamic traveling events were already observed in the vehicle group or at only 1h. However, a significant increase in the relative occurrence of such events as compared to the total number of events was found in the LPS group 48 h post-injection, as compared to the vehicle (+25.5%, p = 0.0066; unpaired t-test; Fig. 2e). The peak amplitude was increased but non-significantly (mean difference +2.2%, U(6,6) = 7, p = 0.0931; Mann–Whitney test; Fig. 2f), while the propagation distance (+2.6 mm, p=0.0313, unpaired t-test; Fig. 2g) of the traveling waves were also significantly increased at 48 h post-injection of LPS.

**Figure 2:**
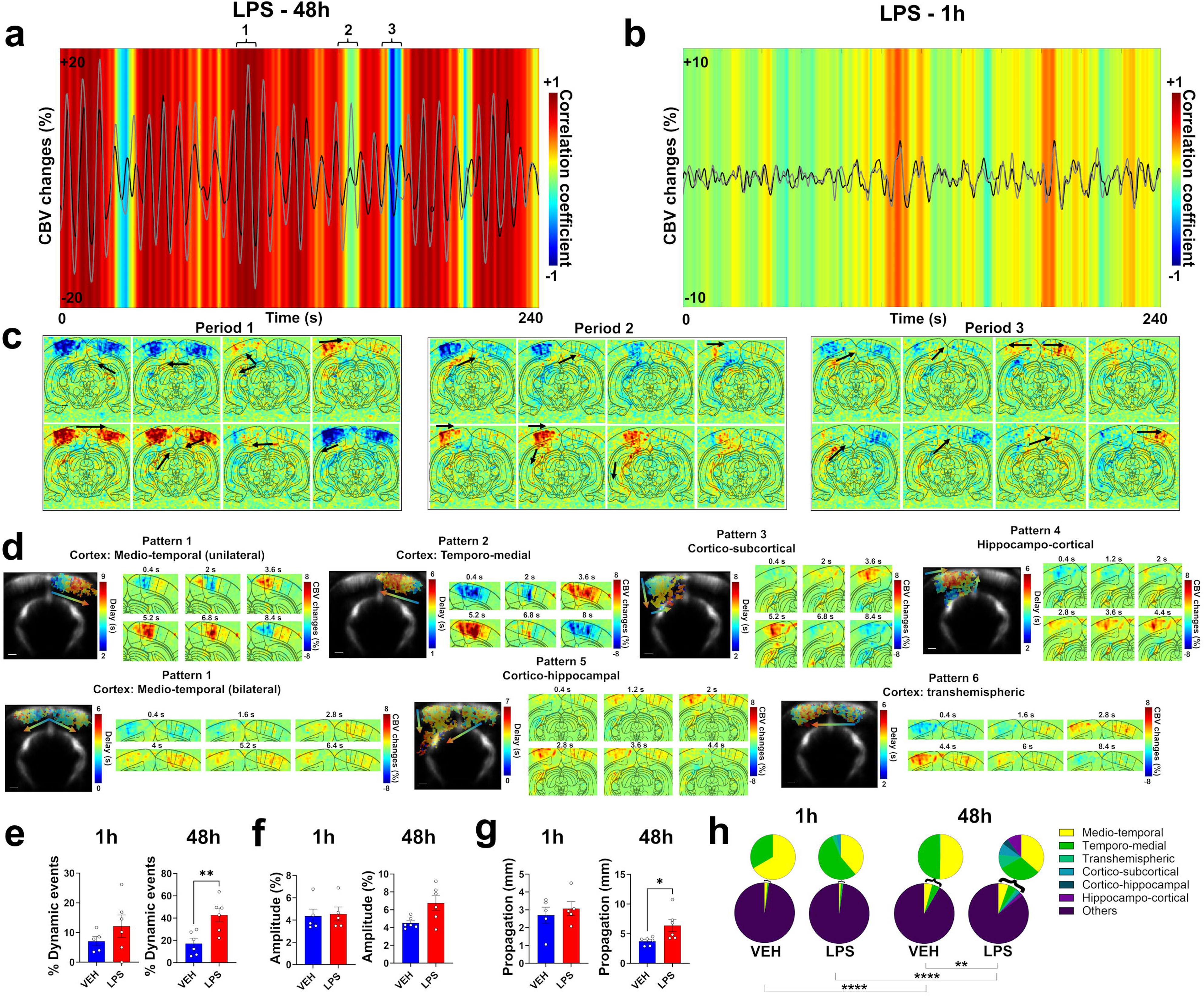
Spatial and temporal dynamics of CBV oscillations. (a) Dynamic functional connectivity analysis in a representative individual, 48h after LPS injection. The same individual is shown in (b), only 1h after LPS injection. The CBV changes over time in the left and right visual cortex are shown in grey and black curves, respectively. The dynamic connectivity between the left and right visual cortex is shown in the colored heatmap, using a sliding window of 6 seconds and a step of 0.4 second. The hemodynamic oscillations are associated with small epochs of hyperconnectivity or anticorrelation between cortical hemispheres. Corresponding changes in CBV, pixel-by-pixel, are mapped during such periods in (c), showing that such changes in dynamic functional connectivity reflect different patterns of hemodynamic traveling waves that can occur almost simultaneously across the bilateral cortex (Period 1 of hyperconnectivity), unilaterally (Period 2 of low connectivity), or sequentially between the two sides (Period 3 of negative connectivity). (d) Different patterns of traveling waves identified across all acquisitions after clustering of peak events, showing that waves could travel along different directions across the cortex and also propagate from subcortical areas to the cortex, and reversely. Examples of delay maps and corresponding maps of CBV changes across time are shown for each pattern. (e) Percentage of dynamic traveling events identified as compared to the total number of events including static and localized increases of CBV, showing that traveling waves were observed more often in the LPS group 48h after injection. Each bar is the mean across individuals ± SEM. n=5 for VEH-1h, n=6 for other groups. (f) Amplitude of the CBV increases for the traveling events, showing that the traveling waves were of higher amplitude in the LPS group 48h after injection. Each individual point is the mean for all dynamic events detected over the acquisition. Each bar is the mean across individual points ± SEM. n=5 for VEH-1h and LPS-1h, n=6 for other groups. (g) Propagation distance for the traveling waves, showing that the waves traveled on longer distances in the LPS-48h group. Each individual point is the mean for all dynamic events detected over the acquisition. Each bar is the mean across individual points ± SEM. n=5 for VEH-1h and LPS-1h, n=6 for other groups. Unpaired t-tests in (e), (f), (g), except a Mann-Whitney test for 48h condition in (f) due to variance heterogeneity. (h) Pie charts showing the proportion of each traveling pattern as compared to the total number of detected events. To better visualize the relative occurrence of each pattern, the proportions are also shown with inlays excluding the unclassified events. Two-way ANOVAs for between-groups comparisons and mixed-effect analyzes for between-time points comparisons. n=5 for VEH-1h, n=6 for other groups.

As we individually classified each event, we studied the frequency of occurrence of events depending on their pattern of propagation (Fig. 2h). The medio-temporal or temporo-medial waves, either in one hemisphere or bilaterally, were found in LPS and VEH groups at both time points. However, their occurrence was always higher at 48 h as compared to 1 h for all groups, leading to significant differences in terms of global patterns representation. The distribution of the patterns was also significantly different between LPS and VEH at 48 h, with an increase in the relative proportion of cortical events and the appearance of propagations between cortical and subcortical areas in both directions (VEH: 4.2%, 4.1%, 0.1% and 91.6% of medio-temporal, temporo-medial, transhemispheric and unclassified/static events, respectively; LPS: 5.3%, 4.5%, 1.4%, 1.2%, 0.7%, 1.5% and 85.4% of medio-temporal, temporo-medial, transhemispheric, cortico-subcortical, cortico-hippocampal, hippocampo-cortical and unclassified/static events, respectively; F_(6,70)_ = 4.2, p = 0.0012, interaction between patterns factor and group factor).

To summarize, the hemodynamic oscillations around 0.1 Hz displayed propagational properties, reflected by an increase of occurrence of traveling waves in the LPS group that were also of higher amplitude and longer distance of propagation. They were mainly located in the cortex, not only in pial vessels but largely involving vertical penetrating vessels. They also occasionally involved the hippocampus or other subcortical regions, but only in the LPS group.

### Reconfiguration of resting hemodynamic activity towards oscillatory modes following inflammation

To further characterize the contribution of these oscillations on the global hemodynamic fluctuations, a principal component analysis (PCA) was performed on the mean covariance matrix for each group (see Methods). Compared to the VEH group (Fig. 3a-b) and the immediate effects after 1 h (Fig. 3c), the injection of LPS alone produced at 48 h spatial maps of principal components with highly fragmented cortical regions, characterized by small, complex patches with correlated or anticorrelated activity, indicating a fine-grained functional subdivision of the cortex (Fig. 3d). The frequency spectra of the temporal signature of the components for this specific condition showed frequency peaks between 0.08 and 0.15 Hz (Fig. 3d), as observed previously with the raw Power Doppler signal in these regions. Strikingly, at 48 h post-injection, the first eight components alone were sufficient to explain more than 99.5% of the variance for the LPS group, as opposed to the VEH group or to the early time-point in which more than 1000 components were required (Fig. 3e), highlighting the importance of this phenomenon. For easier comparison, the amount of explained variance by the first ten components were 82.3% for VEH-1h, 78.3% for VEH-48h, 88.8% for LPS-1h, and 99.6% for LPS-48h. In other words, the resting state activity under inflammation could be fully described by these large-amplitude hemodynamic oscillations occurring at a handful of distinct spatial nodes with their own frequency spectra.

**Figure 3:**
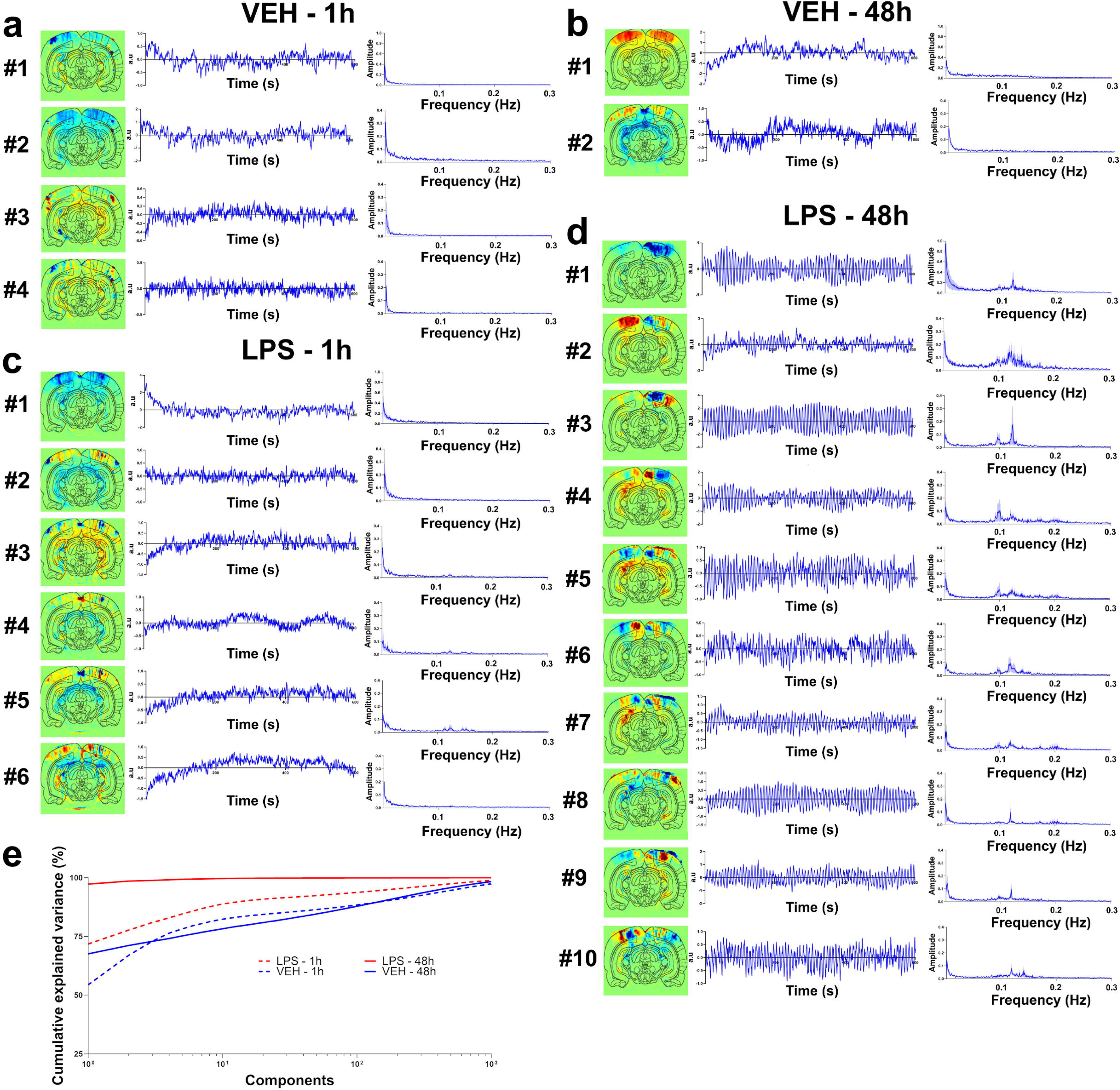
Principal component analysis reveals oscillatory modes following LPS exposure. Spatial, temporal and spectral signature of the principal components in the vehicle group, 1h (a) and 48h (b) after injection, and in the LPS group, 1h (c) or 48h (d) after injection (group analysis on n=6 per group). Only the components with eigenvalues >0.1 are shown. The spatial and spectral signatures are obtained from the group averaging; the temporal profile is shown for representative individuals. Resonant modes can be visualized in the LPS 48h group with peaks in the frequency spectrum between 0.08 Hz and 0.15 Hz, cutting the cortex into distinct subterritories. (e) Scree plot showing the percentage of explained variance for each component. The ten first components in the LPS 48h group can explain more than 99% of the variance, contrary to the other groups.

### Hemodynamic waves in neuroinflammatory conditions are related to the reactivity of glial cells

As LPS-induced inflammation is known to heavily modify the morphology and function of microglia and astrocytes (that become “reactive” in such conditions), we decided to investigate the relationship between glial cells and inflammatory vasomotion. For studying microglial reactivity, we used a new data-driven approach that relies on principal component analysis of multiple morphological features of segmented microglial cells^40^, visualized by Iba-1 immunofluorescence on cerebral slices of the same animals that were used for fUSi. For the astrocytic contribution, we chose a different strategy since astrocytes’ morphology is very complex and usual astrocytic protein markers are not able to label the numerous finest processes that would be needed for assessing the overall cell morphology. Instead, we used additional experimental groups that relied on different intracerebral injections of gliotoxins that are known to induce specific lesions of astrocytes through different mechanisms, namely alpha-amino-adipate (AAA, which acts by blocking the glutamine synthase^42^) and fluorocitrate (FC, which blocks the Krebs cycle after selective uptake in astrocytes^43,44^). These gliotoxins were injected alone or in addition to LPS before fUSi to evaluate the influence of astrocytes on the hemodynamic oscillations, before sacrificing the animals.

Using relevant immunostainings, we first verified the specificity of astrocytic lesions and absence of obvious toxicity on other cell types in those groups, as shown in Supplementary Fig. 1a. The staining intensity of GFAP (an astrocytic protein upregulated in reactive astrocytes) globally differed between groups (H(6,192) = 62.53, p<0.0001; Kruskal-Wallis test, FDR correction; Supplementary Fig. S1b). It was more pronounced in the LPS group compared to the vehicle injection, although non significantly (Mean rank diff = +20.14, q = 0.0789). Conversely, the groups that received the gliotoxin FC alone or in addition to LPS displayed significantly reduced GFAP staining (FC: Mean rank diff = -45.49, q = 0.0008; LPS+FC: Mean rank diff = -55.79, q = 0.0003), whereas the gliotoxin AAA alone did not inhibit GFAP expression significantly (Mean rank diff = -12.63, q = 0.1541), but did when combined with LPS (Mean rank diff = -67.90, q <0.0001). Although GFAP is often used as an astrocytic marker, it is not recommended alone to assess overall astrocytes density, as it is overexpressed in reactive state, and only few astrocytes express GFAP in cortical grey matter. For these reasons, we also performed NDRG2 (a marker of cortical astrocytes^45^) labelling, which also showed variable staining between groups (H(6,192) = 28.22, p <0.0001). All groups receiving either only gliotoxins or a gliotoxin in co-injection with LPS showed significant decrease of NDRG2 intensity (AAA: Mean rank diff = -38.44, q = 0.0012; FC: Mean rank diff = -54.89, q <0.0001; LPS+AAA: Mean rank diff = -33.44, q = 0.0044; LPS+FC: Mean rank diff = -48.55, q = 0.0003; Supplementary Fig. S1b) in comparison to the vehicle group. These observations confirmed an astrocytic lesion at the injection site by AAA or FC. In contrast, no significant modification of NDRG2 level was observed after LPS administration alone, indicating that LPS did not induce an astrocyte loss (Mean rank diff = -3.337; q = 0.1704). Furthermore, despite an overall group effect (H(6,192) = 25.69, p = 0.0001), no significant difference in neuronal density (NeuN+ cells counting) was observed between each group and vehicle (LPS, Mean rank diff = -28.97 , q = 0.1010 ; AAA, Mean rank diff = +20.91, q = 0.2365; FC, Mean rank diff = +7.298, q = 0.6322; LPS+AAA, Mean rank diff = -11.52, q = 0.5549; LPS+FC, Mean rank diff = +34.21, q = 0.1010; Supplementary Fig. S1b), suggesting that the injections did not induce neurotoxicity in our models. Finally, we also compared the expression of the microglial marker Iba-1 between vehicle and other groups (H(6,192) = 12.27, p = 0.0312), that remained unchanged in most groups (LPS, Mean rank diff = +0.4563, q = 0.6136; AAA, Mean rank diff = -17.72, q = 0.184; LPS+FC, Mean rank diff = -14.51, q = 0.2364) but was slightly decreased in the FC and LPS+AAA groups (FC, Mean rank diff = -32.58, q = 0.0203 ; LPS+AAA, Mean rank diff = -35.94, q = 0.0203; Supplementary Fig. S1b), suggesting that global density of microglia was unchanged by LPS (consistently with an expression of Iba-1 in both activated and quiescent microglial cells^46^) but were occasionally slightly lowered by the astrocytic toxins.

Then we focused on the morphology of microglial Iba-1+ cells. As expected, the extensive morphology analysis showed a switch from a ramified to an ameboid morphology 48h following LPS injection (Fig. 4a, segmented cells in red, as compared to the vehicle in blue). Numerous parameters (see ^40^ for the definition of all parameters) were investigated (One-way ANOVAs followed by Dunnett’s corrections, unless otherwise specified) ; the perimeter area ratio (Mean rank diff = -17.75, q = 0.0011, Kruskal-Wallis test) the ramification index (Mean rank diff = -17.75, q = 0.0016, Kruskal-Wallis test), the processes/cell areas ratio (Mean diff = -0.2028, p = 0.0010), the skeleton/processes ratio (Mean diff = -9.329, p = 0.0104), the polarization index (Mean diff = -0.1410, p = 0.0001), the lacunarity (Mean diff = -0.0007, p = 0.0247), the density (Mean diff = -0.0038, p = 0.0162), the branching index (Mean diff = - 0.0909, p = 0.0203), the inertia (Mean diff = -1.8390, p = 0.0107), the processes/soma areas ratio (Mean diff = -1.2920, p = 0.0162), the convexity (Mean diff = -0.3806, p = 0.0324), the convex hull radii ratio (Mean diff = -0.4156, p = 0.0076), the endpoints/branch points ratio (Mean diff = -0.9093, p = 0.0078) and the fractal dimension (Mean diff = -0.0875, p = 0.0241) were significantly decreased 48h after LPS injection as compared to vehicle injection, while the circularity (Mean rank diff = +17.50, q = 0.0020, Kruskal-Wallis test), the roundness factor (Mean diff = +0.7908, p = 0.0004), the convex hull circularity (Mean diff = +0.0746, p = 0.0004), and the solidity (Mean rank diff = +16.65, q = 0.0025, Kruskal-Wallis test) were increased (Fig. 4b and Supplementary Fig. S1c). The features span ratio and linearity showed no significant differences. The other additional groups that received astrocytic toxins were also assessed for microglial morphology (Fig. 4a-b). No significant morphological differences with control condition were observed 48h after the injection of each astrocytic toxin alone. When gliotoxins were co-injected with LPS, morphological differences were still observed compared to vehicle, although they were slightly less important than in the LPS alone group. Indeed, the perimeter area ratio (LPS+AAA, Mean rank diff = -12.25, q = 0.0165 ; LPS+FC, Mean rank diff = -13.50, q = 0.0132; Kruskal-Wallis test), the ramification index (LPS+AAA, Mean rank diff = -12.25, q = 0.0165; LPS+FC, Mean rank diff = 12.50, q = 0.0165; Kruskal-Wallis test), the processes/cell areas ratio (LPS+AAA, Mean diff = -0.1387, p = 0.0340; LPS+FC, Mean diff = -0.1723, p = 0.0073), the skeleton/processes ratio (LPS+AAA, Mean diff = -7.914, p = 0.0448; LPS+FC, Mean diff = -8.569, p = 0.0275), the polarization index (LPS+AAA, Mean diff = -0.0935, p = 0.0119; LPS+FC, Mean diff = -0.1067, p = 0.0041), the circularity (LPS+AAA, Mean rank diff = +11.75, q = 0.0313; LPS+FC, Mean rank diff = +13.50, q = 0.0198; Kruskal-Wallis test), the roundness factor (LPS+AAA, Mean diff = +0.5152, p = 0.0282; LPS+FC, Mean diff = +0.6370, p = 0.0059), and the solidity (LPS+AAA, Mean rank diff = +12.25, q = 0.0165; LPS+FC, Mean rank diff = +13.25, q = 0.0150; Kruskal-Wallis test) were modified in both the LPS+AAA and LPS+FC groups. Other features were significantly different only for the LPS+FC group, such as convex hull circularity (Mean diff = +0.0500, p = 0.0216), branching index (Mean diff = -0.0843, p = 0.0454), endpoints/branch points ratio (Mean diff = -0.8977, p = 0.0129) and processes/soma areas ratio (Mean diff = -1.197, p = 0.0379) (Fig. 4b and Supplementary Fig. S1c). Finally, Andrew’s score ranking, which integrates multiple parameters that characterized best our dataset, was used to classify each microglial cell from the different animals in terms of reactivity (a negative score depicts a round-shape morphology, typical of the reactive state, whereas a positive score depicts a well-ramified, basal morphology – see Fig. 4c for the relative frequency of microglia in each group as a function of their Andrew’s score). The frequency of reactive microglia was much higher in the LPS alone as compared to vehicle and to any other group. A slight increase of microglial reactivity was found in the FC group, while a moderate increase was found for the LPS+gliotoxins groups, as shown by the mean of Andrew’s scores between the groups (Fig. 4d ; F_(5,20)_ = 5.377, p = 0.0027; One-way ANOVA, Dunnett correction).

**Figure 4:**
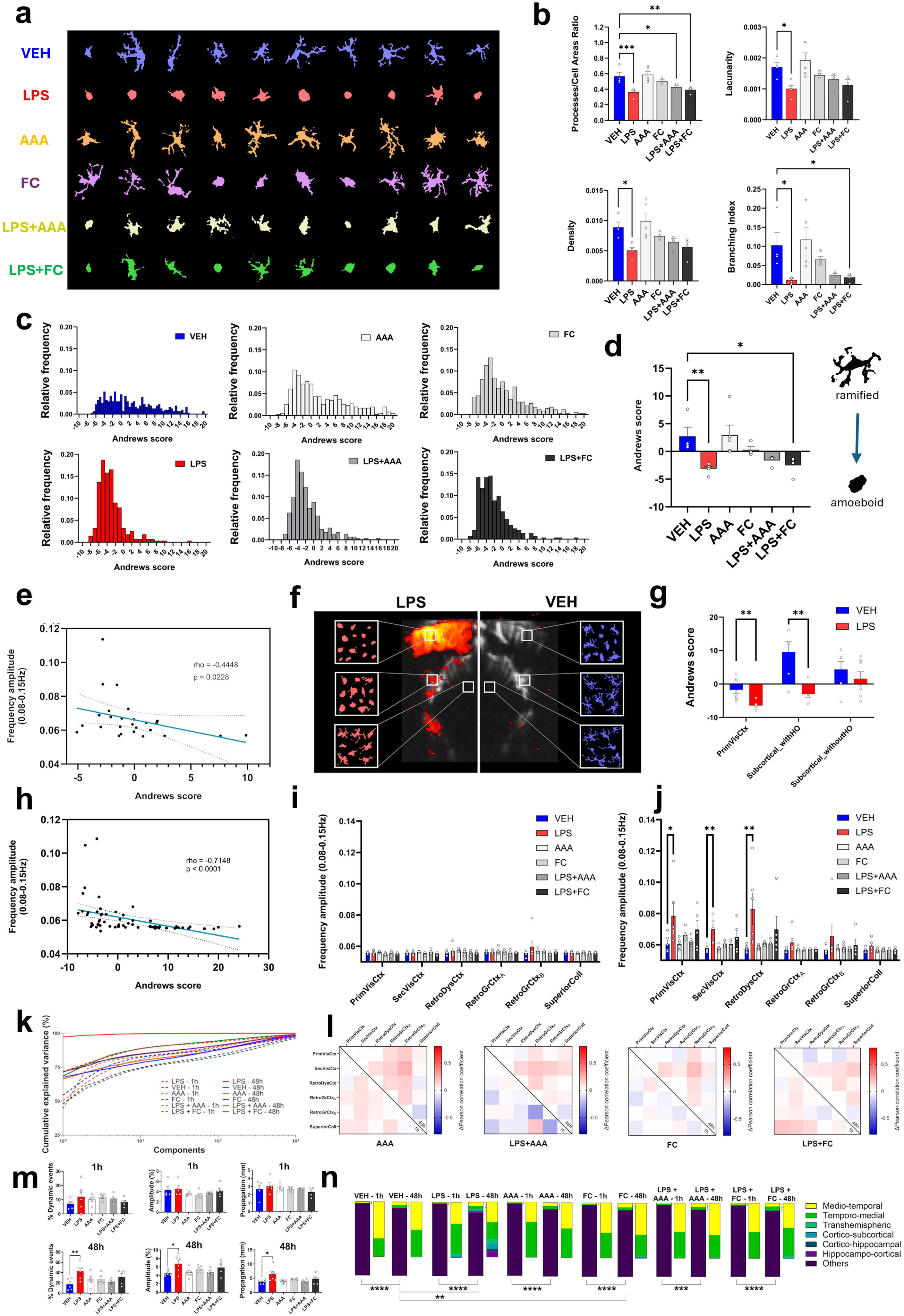
Relationship between reactive glial cells and CBV oscillations. (a) Representative illustrations of segmented microglia for each group (vehicle, LPS, and/or gliotoxins injections). (b) Main parameters that best describe the microglia morphology in the dataset in the different groups. One-way ANOVAs followed by Dunnett’s multiple comparisons test. Each bar is the mean ± SEM. n=5 for LPS and AAA, n=4 for other groups. One point per animal, average of 78.8 ± 3.1, 55.6 ± 28.7, 71 ± 20.8, 86.5 ± 9.4, 71.25 ± 24.4 and 70.5 ± 16.3 microglial cells by animal for VEH, LPS, AAA, FC, LPS+AAA and LPS+FC, respectively. (c) Distribution graph of Andrew’s scores of microglial cells (that estimates the reactivity of microglia based on their morphology) in the different groups. (d) Bar plot of the global Andrew’s score for each group. One-way ANOVA followed by Dunnett’s multiple comparisons test. Each bar is the mean ± SEM. (e) Scatter plot and linear regression showing the relationship between CBV oscillations in 0.08-0.15 Hz range and global Andrew’s score in individuals from all groups (n=26). (f) Illustration of the spatial correlation between microglial morphology and hemodynamic waves. Individual maps of fALFF are shown for a representative individual from LPS group (left) and VEH group (right), with corresponding areas where microglia were cropped. From top to bottom: primary visual cortex, subcortical area with oscillations, and subcortical area without oscillations. (g) Bar plot of the global Andrew’s score for the three sets of regions. Multiple Mann-Whitney tests with FDR correction of 5%. Each bar is the mean ± SEM. VEH: n=3 to 4 depending on regions, one point per hemisphere, average of 37.0 ± 7.3 microglial cells by animal and region hemisphere; LPS: n=5, one point per hemisphere, average of 32.2 ± 11.3 microglial cells by animal and region hemisphere. (h) Scatter plot and linear regression showing the relationship between CBV oscillations in 0.08-0.15 Hz range and global Andrew’s score among the three sets of regions of interest (primary visual cortex/subcortical with oscillations/subcortical without oscillations, total of 66 points) for individuals of the VEH and LPS groups (10 animals). (i) Frequency amplitude of CBV changes in 0.08-0.15 Hz range at 1 h post-injection for all experimental groups. GLM followed by Dunnett’s correction. n=6 for VEH and LPS, n=4 for LPS+AAA, n=5 for other groups. (j) Frequency amplitude in 0.08-0.15 Hz range at 48 h post-injection for all experimental groups. GLM followed by Dunnett’s correction. Each bar is the mean ± SEM. n=6 for VEH, LPS and AAA, n=4 for LPS+AAA, n=5 for other groups. (k) Scree plot showing the percentage of explained variance for each component after PCA for all groups. The ten first components in the LPS 48h group can explain more than 99% of the variance, contrary to all the other groups that received gliotoxins alone or coinjected with LPS. (l) Functional connectivity changes after 1 and 48 hours. Matrices of the correlation coefficient variations of each group (that received gliotoxin alone or in combination with LPS) compared to vehicle are shown (bottom-left for 1 h post-injection, top-right for 48 h post-injection). For statistical comparisons, a Fisher Z-transformation was applied to the correlation coefficients before two-way ANOVA followed by Dunnett’s multiple comparisons test (all non-significant). n=6 for VEH and LPS, n=4 for LPS+AAA, n=5 for other groups at 1h, n=6 for VEH, LPS and AAA, n=4 for LPS+AAA, n=5 for other groups at 48h. (m) Percentage, amplitude and propagation of traveling waves for all groups at 1h and 48h post-injection. Each bar is the mean across individuals ± SEM. One-way ANOVAs followed by Dunnett’s tests. n=6 for LPS (or n=5 for amplitude and propagation due to no traveling event in one animal), n=4 for LPS+AAA, n=5 for other groups at 1h; n=6 for VEH, LPS and AAA, n=4 for LPS+AAA, n=5 for other groups at 48h. (n) Bar charts showing for all groups the proportion of each traveling pattern as compared to the total number of detected events. To better visualize the relative occurrence of each pattern, the proportions are also shown excluding the unclassified events. Two-way ANOVAs for between-groups comparisons and mixed-effect analyzes for between-time points comparisons. n=6 for LPS, n=4 for LPS+AAA, n=5 for other groups at 1h; n=6 for VEH, LPS and AAA, n=4 for LPS+AAA, n=5 for other groups at 48h.

As fUSi was performed for all animals of these different groups, the microglial morphological changes could be correlated with the amplitude of hemodynamic oscillations for all animals that received the different injections. Interestingly, the Andrew’s score of overall cell morphology was negatively correlated with the increase of the frequency amplitude of 0.08-0.15 Hz range in the same animals (Fig. 4e; rho: -0.445; p = 0.023; Spearman correlation test). In other words, as negative values of Andrew’s score are reflective of round-shaped morphology, the rats that displayed the most reactive microglial cells showed higher amplitudes of vasomotion. Significant correlations were also observed for many individual morphological parameters in addition to the global Andrew’s score (results not shown). This finding raised the important possibility of a spatial correlation between hemodynamic oscillations and Andrew’s score, as we previously identified that hemodynamic waves also extend outside the injected area. Accordingly, we further investigated the morphology of microglial cells in the LPS and VEH groups at 48h post-injection, in three different sets of regions depending on the spatial patterns of inflammatory waves (from top to bottom, Fig. 4f): the injected area (primary visual cortex), subcortical regions with oscillations (from the hippocampus and dorsal midbrain), and subcortical regions without oscillations (from the ventral midbrain). We found a significantly lower Andrew’s score in LPS compared to VEH (Fig. 4g), not only in the primary visual cortex (Mean rank diff = -6.964, q = 0.001224; Mann–Whitney tests, FDR correction) but also in the distant regions displaying hemodynamic oscillations (Mean rank diff = -6.190, q = 0.001224), indicating the presence of reactive microglia in these regions and suggesting that inflammation in our model can spread outside the injection site. In contrast, in the regions that were devoid of oscillations in the LPS group, we found no differences of microglial morphology between groups (Mean rank diff = -1.857, q =0.155828). Then, we evaluated the global relationship between Andrew’s score and the frequency amplitude in the 0.08-0.15 Hz range from the same regions by pooling all measurements from the three different sets of regions, including both VEH and LPS groups. There was a highly significant, negative correlation between fALFF and Andrew’s score (Fig. 4h; rho= -0.715, p < 0.0001; Spearman correlation test), suggesting that the amplitude of hemodynamic oscillations correlates not only to the overall changes in glial reactivity in each individual, but also to the spatial extent of the glial reactivity, and that the local presence of reactive glia is needed for this vasomotion amplification to occur.

Focusing back on the effect of gliotoxins for further investigation of the causal relationship between reactive glia and hemodynamic oscillations, their injection alone did not significantly impact the amplitude of vasomotion compared to the control animals in any region at both time points (Fig. 4i-j), but the LPS-induced increase of amplitude at 48h (Primary visual cortex: Estimate = +3.81, p = 0.0165; Secondary visual cortex: Estimate = +2.98, p = 0.00164; Retrosplenial dysgranular cortex: Estimate = +5.32, p = 0.00456; GLM with Dunnett’s correction) was fully prevented by AAA and reduced by FC co-injection (Primary visual cortex: LPS+AAA, Estimate = +0.459, p = 0.986; LPS+FC, Estimate = +2.22, p = 0.333; Secondary visual cortex: LPS+AAA, Estimate = +0.756, p = 0.806; LPS+FC, Estimate = +1.93, p =0.0814; Retrosplenial dysgranular cortex: LPS+AAA, Estimate = +0.933, p = 0.947; LPS+FC, Estimate = +3.06 p = 0.231). This suggests that the inflammatory vasomotion in the LPS group alone depends on the local presence of astrocytes. Accordingly, when comparing the contributions of hemodynamic oscillations to the overall signal changes in LPS with the other groups that received the astrocytic toxins using PCA, here again only the variance from the LPS-48h group was fully described by less than 10 components, whereas FC or AAA alone or combined with LPS required about 1000 components to explain the same amount of variance, similarly to the VEH group (Fig. 4k). For comparison, the amount of explained variance by the 10 first components was 80.6% for AAA-1h, 79.8% for AAA-48h, 74.6% for FC-1h, 84.8% for FC-48h, 82.8% for LPS+AAA-1h, 84.4% for LPS+AAA-48h, 73.9% for LPS+FC-1h, and 88.6% for LPS+FC 48h (whereas this amount was 99.6% for LPS-48h). Moreover, no more increase of resting-state functional connectivity was found at 48h post-injection of LPS when it was co-injected with AAA or FC, while none of these molecules alone produced any effect on functional connectivity at 1h or 48h post-injection (Fig. 4l). In parallel, the spatial maps of the components from the different gliotoxins groups (see Supplementary Fig. S2) displayed broader, more globally coherent activity patterns than LPS alone, with entire cortical areas showing correlation or anticorrelation with subcortical structures, and occasionally, inter-hemispheric anticorrelation (i.e. component 3 for the FC group at 1 h and 48 h – Supplementary Fig. S2c-d). Interestingly, the injection of gliotoxins alone or with LPS tended to produce acute subcortical effects, particularly in the hippocampus, that are visible on the spatial maps of the components at 1 h post-injection (i.e. components 3 for AAA, 2 for FC, 1 for LPS+AAA and 2 for LPS+FC – Supplementary Fig. S2a, c, e and g, respectively). In terms of frequency properties of the components, only certain components of the LPS+FC group still showed frequency peaks around 0.1 Hz at 48 h. We also noticed that the gliotoxins tended to produce irregular very low frequency fluctuations at the acute phase, 1 h post-injection, as showed in the temporal signature of some components in Figure S2 (i.e., components 3 to 5 of AAA, 2 to 3 of FC, and 1 and 2 of LPS+FC). Taken together, these PCA results show that LPS oscillations at 48h were respectively attenuated or fully blocked by FC and AAA, whereas these toxins alone produced distinct hemodynamic effects by themselves that were most obvious at 1h.

When analyzing the properties of hemodynamic changes as traveling waves, it also appeared that LPS combined with AAA or FC was not associated to any significant change in the proportion (LPS+AAA, +4.0%, p = 0.9833, LPS+FC, +14.1%, p = 0.2480; One-way ANOVA followed by Dunnett’s correction), peak amplitude (LPS+AAA, +0.2%, p = 0.9987, LPS+FC, +1.4%, p = 0.3411) or propagation (LPS+AAA, +0.1 mm, p > 0.9999, LPS+FC, +0.9 mm, p = 0.7374) of such waves (Fig. 4m), with no effect of the gliotoxins alone. The combination of AAA or FC with LPS was also found to restore the normal profile of traveling waves spatial patterns (Fig. 4n), that is, mostly medio-temporal or temporo-medial waves. We still observed an overall difference of waves repartition with less unclassified events for all groups at 48h compared to 1h, and FC alone was significantly different from VEH at 48h due to the appearance of some waves involving the subcortical structures or transhemispheric propagation (5.6%, 3.7%, 0.3%, 0.06%, 0.06%, 0.06% and 90.2% of medio-temporal, temporo-medial, transhemispheric, cortico-subcortical, cortico-hippocampal, hippocampo-cortical and unclassified/static events, respectively; F_(6,63)_ = 2.624, p = 0.0247).

To summarize, the injection of toxins that induce lesions of astrocytes, in combination with LPS, was followed by an almost complete normalization of the inflammatory vasomotion, as shown by the different parameters that we assessed (amplitude of oscillations, functional connectivity, traveling waves proportion and trajectories, and variance explained by principal components). Moreover, when taking into account all animals from the different groups, a correlation was found between the microglial morphology changes and the amplitude of vasomotion. The spatial extent of oscillations also appeared to be correlated with the areas displaying microglial changes in the LPS model. Altogether, this suggest that microglia and astrocytes both participate (and most probably, interact together) in this hemodynamic phenomenon.

### Kinetics of hemodynamic waves induction after LPS exposure

To understand the kinetics of the oscillatory effect resulting from LPS injection, the amplitude of 0.08-0.15 Hz frequencies was evaluated at 1- and 7-days post injection in addition to the 1h (0 day) and 48h (2-days) groups (Fig. 5a). The animals without injection (wI; only with a skull thinning) did not differ from animals measured 1 h after LPS injection (Estimates ranging from +0.14 to +1.93 and p values from 0.999 to 0.423 depending on regions; Generalized Linear Model, Dunnett correction). As expected, 48 h after injection, the amplitude of the frequency range of interest was increased in the primary visual cortex (Estimate = +4.84, p = 0.00429), the secondary visual cortex (Estimate = +3.40, p = 0.00705) and the retrosplenial dysgranular cortex (Estimate = +5.43, p = 0.000847), compared to 1 h post-injection. The increase was already present in the primary and secondary visual cortices, 24 h after the intracerebral injection of LPS (primary visual cortex: 24 h, Estimate = +5.95, p = 0.00214; secondary visual cortex: 24h, Estimate = +5.01, p = 0.000640), and was no longer different from baseline state 7 days after injection (primary visual cortex: 7 d, Estimate = +1.10, p = 0.893; secondary visual cortex: 7 d, Estimate = +0.704, p = 0.919 ; retrosplenial dysgranular cortex: 7 d, Estimate = +1.01, p = 0.905). In this longitudinal analysis, a significantly higher fALFF was also found in other regions, such as the retrosplenial granular cortex A at 48h (Estimate = +1.70, p = 0.0289) and the superficial part of the superior colliculus at 24h (Estimate = +1.53, p = 0.00605) and 48h (Estimate = +1.19, p = 0.00833). Therefore, the appearance of hemodynamic oscillations was present in the earlier phase of neuroinflammation and was no more observed after one week.

**Figure 5:**
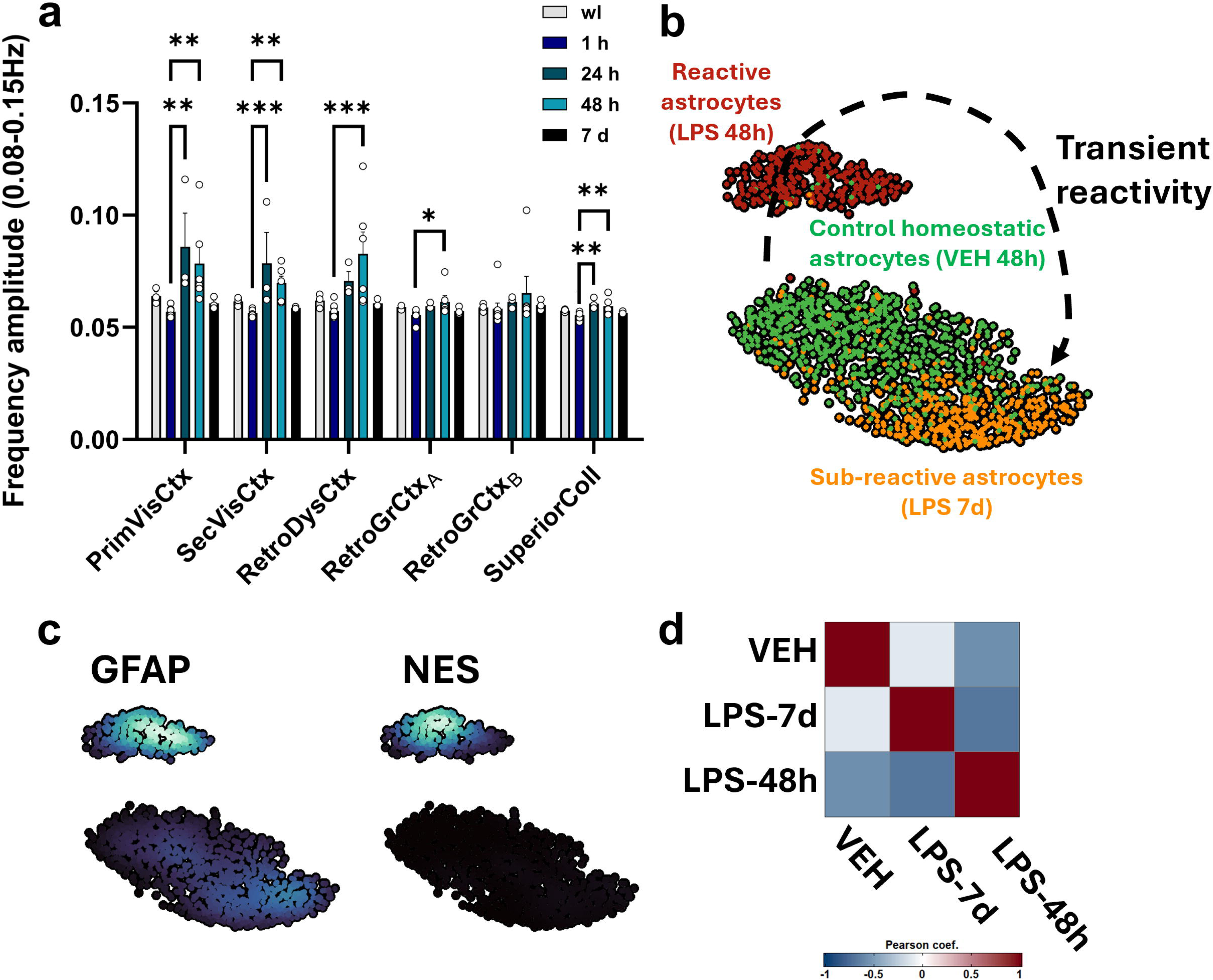
Kinetics of LPS effects on the hemodynamic oscillations and astrocytes reactivity. (a) Frequency amplitude in 0.08-0.15 Hz range in animals without any injection (wI), or at 1h, 24h, 48h (2d) or seven days (7d) after injection. GLM followed by Dunnett’s correction. Each bar is the mean ± SEM. n=9 at 1h, n=6 at 48h, n=3 at other time points. The oscillations were already present at 24h post-injection but disappeared after 7 days. (b) UMAP plot showing the distribution of astrocytes in the three conditions: 48h post-injection of VEH in green or LPS in red, and 7 days post-injection of LPS in yellow. Note that at 7 days post-injection of LPS, the molecular phenotype of astrocytes is much closer to the VEH group than at 48h post-injection. RNA-seq performed on n=2 rats per group. (c) Density plot showing the selective increase of GFAP and Nestin expression in the cluster that corresponds to LPS at 48h post-injection, but not in the cluster that corresponds to VEH and LPS-7d groups. (d) Correlation matrix showing the global correlation of gene expression between the different conditions, with higher correlation between LPS-7d and VEH than LPS-48h and VEH. LPS-48h was also markedly different from LPS-7d in terms of global gene expression.

To gain an in-depth overview of the temporality of the intracerebral LPS model at the cellular level, we performed large-scale single-nuclei profiling from the microdissected visual cortex sampled at 2 days or 7 days following LPS injection, or at 2 days following vehicle injection. Following quality controls, we obtained the transcriptome of 64 471 nuclei by snRNA-seq using PARSE Biosciences split-seq protocol allowing multiplexing. Two brain samples from different individuals were used for each condition and these duplicates were analyzed independently for each condition (Supplementary Fig. S3). Clustering revealed 5 distinct clusters which corresponded to the brain major cell classes (Suppl Fig. S3a). Cell cluster identification was based on the detection of landmark cell type markers, such as *Rbfox3* for neurons, *Aqp4* for astrocytes, *Sox10* for oligodendroglial lineage cells (OLs), *Ptprc* for immune cells and *Cdh5* for vascular cells. The three conditions grouped well together at the general population level (Suppl Fig. S3b) with overall similar proportions of cells but respectively lower and higher relative frequencies of neurons and immune cells at 7d (Suppl Fig. S3c). However, *de novo* clustering of only astrocytes resulted in two different clusters of reactive and homeostatic astrocytes, exclusively enriched in LPS astrocytes at 48h *versus* control VEH astrocytes and LPS astrocytes at 7d, respectively (Fig. 5b,c). Reactive astrocytes over-expressed *Gfap* and *Nes* (Fig. 5c) and genes enriched in wound healing, gliogenesis, glial cell differentiation and response to axon injury biological processes determined using gene ontology (GO) terms enrichment analysis (Suppl. Fig S3d), as compared to vehicle. These enrichments were specific to the reactive astrocytes at 48h and were no more found at 7d, consistently with the UMAP showing that the LPS-7d astrocytes cluster was located much closer to the control cluster, suggesting a peak of reactivity around two days and a return to a state closer to basal astrocytes (Fig. 5b). The direct comparison of astrocytes at 48h post-LPS versus 7 days showed the significant enrichment of the same biological processes (wound healing, gliogenesis, glial cell differentiation and response to axon injury biological processes) as with vehicle injection. The expression of the 2 markers that best described this transient activation at 48h (*Gfap* and *Nes*) went back to normal at 7 days (Fig. 5c). Correlation heatmap between samples confirmed that the whole transcriptional signature of astrocytes from the VEH group correlates more with LPS-7d astrocytes than LPS-48h astrocytes (Fig. 5d). We observed a similar kinetic pattern in Aif1-expressing (Iba1L) microglia, although reactive microglia persisted at LPS-7d (Suppl. Fig. S3e). However, further investigation is needed due to the low number of identified microglial nuclei (as compared to the global number of immune cells, with extensive recruitment of peripheral macrophages at LPS-48h).

### Sustained neurovascular coupling in presence of oscillations

To gain further insights into the functional properties of hemodynamic changes under inflammation, we also evaluated the functional hyperemia changes during visual stimuli (Fig. 6a), which were performed immediately after the resting-state session at both time points (Fig. 1a). As expected in this coronal plane, typical increases of CBV were found in the primary visual cortex and the superior colliculus during the light flickering (Fig. 6b-c). At both 1 h (Fig. 6d-f) and 48 h post-injection (Fig. 6g-i), the response to visual stimuli showed globally no difference between VEH and LPS in the different visual areas (1h: Mean diff ranging from -0.29 to +2.146% and p values from 0.9999 to 0.1711 depending on regions, 2-way ANOVA with Sidak’s correction; 48h: Mean rank differences from -3 to +1 and q values > 0.9999, Mann-Whitney tests with FDR correction). Diving into temporal properties, the time-course of the response at 1h after injection was virtually unchanged compared to vehicle (Fig; 6e), but the oscillations that we observed during resting-state persisted during the visual stimulation at 48h post-injection (Fig. 6h). Interestingly, despite these oscillations, the mean amplitude of the response and the rising and decaying slopes of the curve appeared similar to the vehicle injection, suggesting a sustained neurovascular coupling in this particular state. In fact, in the LPS group specifically, the amplitude of the visual response tended to be higher in animals that displayed the strongest basal oscillations, (as shown in Fig. 6j and in Supplementary Fig. S4a), and there was a significant positive correlation between the fluctuations at 0.08-0.15 Hz during resting-state and the mean amplitude of the functional hyperemia (rho = 0.6084, p = 0.0399; Spearman correlation test - Fig. 6k). This suggests that inflammatory vasomotion shares some common but non-competitive molecular mechanisms with functional hyperemia.

**Figure 6:**
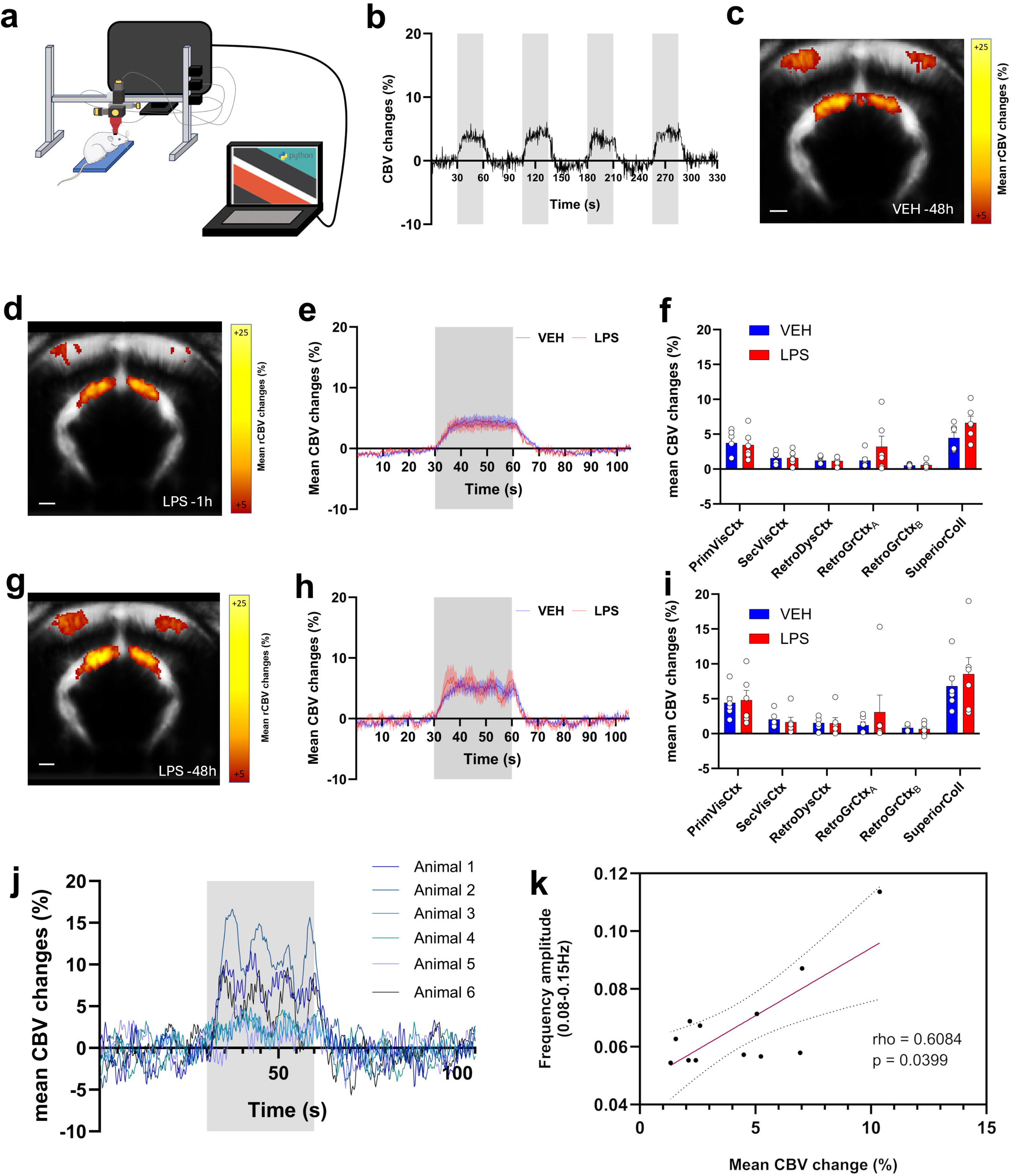
Inflammatory spontaneous oscillations of CBV are associated with increased functional hyperemia during visual stimulations in anesthetized rats. (a) Schematic representation of the experimental setup during fUSi with visual stimulation. (b) Representative time curve of CBV changes in the primary visual cortex of rats following visual stimuli. (c) Mean map of relative CBV changes pixel-by-pixel in the VEH group 48 h post injection. The CBV changes during stimulation after 1h post-injection of LPS in shown as mean map of relative changes in (d), mean time curve in the primary visual cortex in (e) and global changes in the different regions of interest (mean ± SEM; Two-way ANOVA followed by Sidak’s multiple comparisons correction) in (f). n=4 for LPS+AAA, n=5 for FC and LPS+FC, n=6 for other groups. The same results are shown in (g), (h) and (i) for 48h postinjection of LPS, with Mann-Whitney tests with FDR 5% correction in (i) due to non-normality. n=4 for LPS+AAA, n=5 for FC and LPS+FC, n=6 for other groups. (j) Individual mean time curves (averaged across the 4 blocks) for all animals in the LPS group, 48h after injection. (k) Scatter plot and linear regression showing the relationship between amplitude of oscillations during resting-state and the mean CBV changes during visual stimulation in the LPS group at 1h and 48h post-injection (leading to n=12 points).

We also studied the functional hyperemia in the groups that received AAA and FC (Supplementary Fig. S4b-e). The gliotoxins alone had some effects at 1h: AAA led to a modest increase of the visual response in the primary visual cortex at 1h (some time points were significantly higher compared to vehicle from 21 to 30.2 s after the beginning of the stimulation period; Supplementary Fig. S4b), and a global higher response in the superior colliculus (VEH, 4.47% ± 1.85; AAA, 9.88% ± 6.02, p = 0.0012; Supplementary Fig. S4d). For the FC group, the response shape in primary visual cortex was dramatically changed 1 h after the injection, from the usual “plateau”-shape to a dome shape with a much higher amplitude of the response (from 21.2 to 33.2 s after the beginning of the stimulation period; Suppl Fig. S4b). This modification of the response shape was also present after the co-injection of LPS+FC. No obvious effects of gliotoxins, either alone or combined with LPS, was observed when compared to VEH at 48H (Suppl Fig. S4c, e). A slight decrease of visual responses was observed in the primary visual cortex after co-injection of LPS+AAA (from 23.8 to 25 s after the beginning of the stimulation period; Suppl Fig. S4c). To summarize, both gliotoxins acutely impacted the response to visual stimuli in rats, but with different features, notably a strong perturbation of the temporal response of visual cortex following FC injection. In agreement with the previous observations in resting-state conditions, the co-injection of the gliotoxins with LPS prevented the appearance of the vascular oscillations during the visual stimulation 48 h post-injection (as seen in Suppl Fig. S4c,e).

### Hemodynamic waves are not associated to a change in blood flow velocity in cortical vessels

As the hemodynamic oscillations may be related to a global change in vessel tone, we also evaluated quantitative hemodynamic parameters to estimate the level of cerebral perfusion in the vessels using Ultrasound Localization Microscopy (ULM) sequences performed in the same animals (Suppl. Fig. S5), 48 h after intracerebral injections, immediately after the second fUSi session (Fig. 1a). In addition to the density of bubbles (Suppl. Fig. S5a), their absolute velocity (Suppl. Fig. S5b) and their direction (Suppl. Fig. S5c) can be estimated, enabling to identify different vascular compartments (pial vessels, arterioles and venules) in the cortex (Suppl. Fig. S5d). The microbubbles density and velocity were globally unchanged in the different compartments following LPS injection, although the density tended to be higher in arterioles (Velocity: Mean rank diff from -0.9167 in penetrating arterioles to +1.833 in penetrating veinules, all q values > 0.9999; Density: Mean rank diff from +3.117 in pial vessels to +6.05 in arterioles, q from 0.297 in pial vessels to 0.0752 in arterioles - Suppl. Fig. S5e-f), which suggest a slight decrease of the vascular tone. Altogether, this enables to conclude that the appearance of vascular oscillatory waves was not due to a major change of the cerebral blood flow absolute velocity or microbubbles density.

### Confirmation of the LPS-related hemodynamic oscillations in awake rats

As anesthesia could potentially interact with LPS effects on hemodynamic activity, additional rats were imaged in awake freely moving condition using skull implants for probe fitting after progressive habituation (Fig. 7a). For each acquisition, a continuous temporal window of 120 sec was selected based on the absence of any movement artefact (Fig. 7b). Like in anesthetized animals, awake rats showed an increased functional connectivity 48 h after LPS injection between different visual areas (Fig. 7c - primary and secondary visual cortices: Δz-score = 1.126, p = 0.0083; primary visual and retrosplenial dysgranular cortices: Δz-score = 0.8781, p = 0.0448; secondary visual and retrosplenial dysgranular cortices: Δz-score = 0.9915, p = 0.0215). As previously, the PCA showed a reconfiguration after 48 h post-LPS with a very low number of components that could explain the signal variance, as opposed to the baseline measurements or after 1 h (Fig. 7d). Furthermore, the frequency amplitude related to the oscillations were also significantly increased after 48 h compared to the baseline or 1 h post injection in the same regions (Fig. 7e), namely in the primary visual cortex (Baseline: 0.064; 1 h: 0.062; 48 h: 0.080, p = 0.0015), secondary visual cortex (Baseline: 0.061; 1 h: 0.061; 48 h: 0.078, p = 0.0006) and retrosplenial dysgranular cortex (Baseline: 0.064; 1 h, 0.061; 48 h, 0.076, p = 0.0088). An example of long-distance traveling waves in an awake rat can be visualized in Fig. 7f and Supplementary Movie 2. As observed in anesthetized animals, the waves occurred mainly among cortical areas (Wave 1 of Fig.7f is an example of mediotemporal bilateral wave, while Wave 2 followed a cortico-subcortical and bilateral pattern).

**Figure 7:**
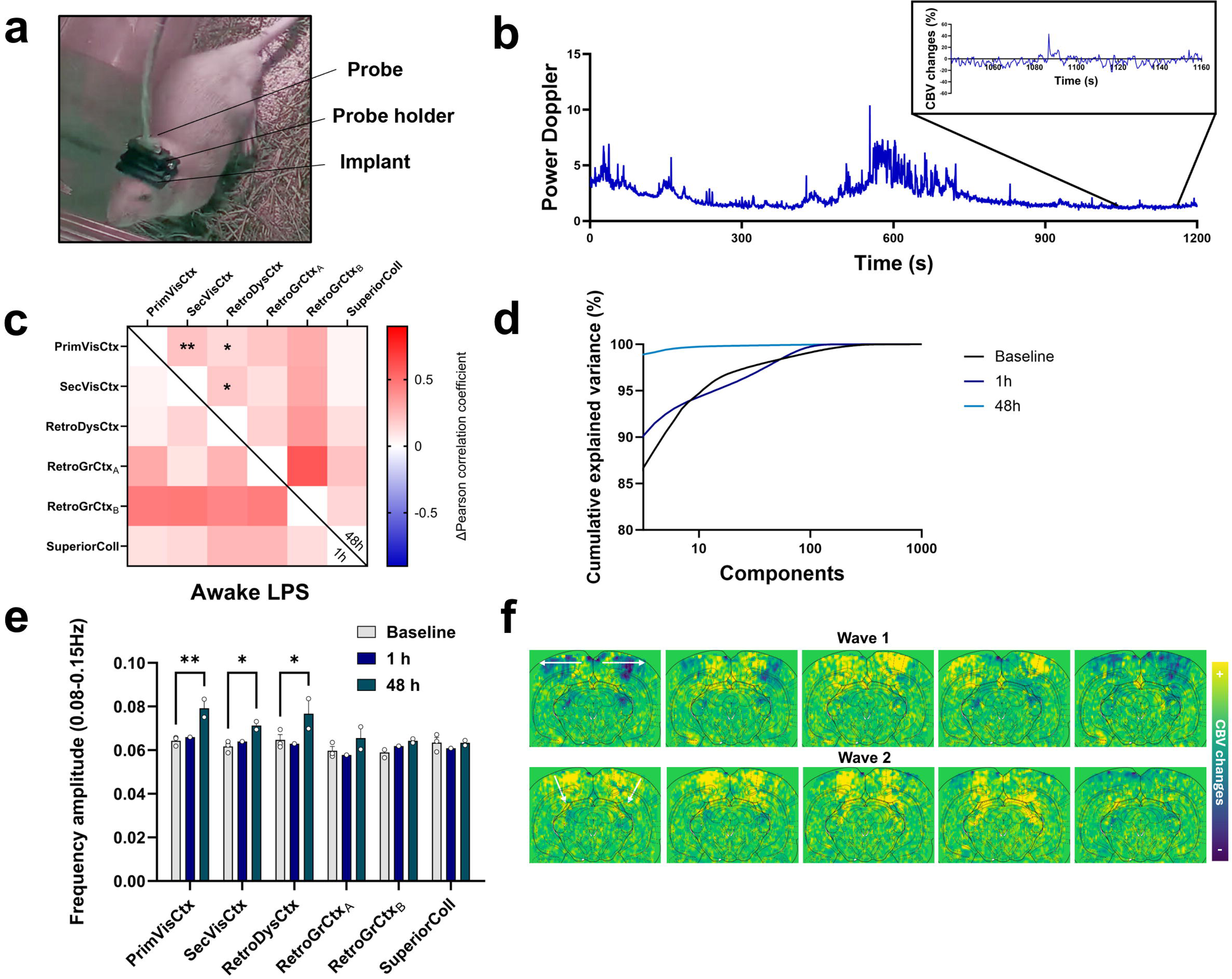
Resting-state fUSi in freely-moving animals revealing hemodynamic oscillations in awake rats after LPS exposure. (a) Illustration showing the probe, probe holder and implant enabling fUSi in freely-moving rats. (b) Representative time-course of the Power Doppler signal in one animal over the 1200s period, enabling to visualize alternance between active periods leading to movement artifacts and quiet state periods, with analysis performed on the consecutive minutes devoid of movement artifact (as shown in the inlay). (c) Functional connectivity matrices 1h or 48h after LPS injection in awake rats. Two-way ANOVA followed by Dunnett’s test. n=3 at baseline, n=1 at 1h, n=2 at 48h. Similarly to anesthetized rats, a significant increase of cortical connectivity was observed at 48h. (d) Scree plot showing the percentage of explained variance for each component obtained after principal component analysis of the different groups (Baseline = without injection, or 1h/48h following LPS injection). Similarly to anesthetized rats, LPS was followed by a global reconfiguration toward oscillatory modes, with less than 10 components that were able to explain almost all the variance in the 48h group only. (e) Frequency amplitude in 0.08-0.15 Hz in awake animals, showing a significant increase of amplitude 48h after LPS injection. Two-way ANOVA followed by Dunnett’s multiple comparisons test. Each bar is the mean ± SEM. n=3 at baseline, n=1 at 1h, n=2 at 48h. (f) Illustrative example of successive maps of relative CBV changes compared to the whole acquisition, (interval 0.8s), 48h post-injection. Two examples of patterns are shown: the medio-temporal wave was the most frequent (Wave 1, top), and other trajectories were occasionally observed involving subcortical areas (Wave 2, bottom). Same acquisition as in Supplementary movie 2 with different colormap and superimposed Paxinos atlas for improved visualization.

## Discussion

While cerebral hemodynamics have been extensively used as a correlate of neuronal activity in animals and humans, they fundamentally arise from complex interactions between many different cellular components. Indeed, neuronal, glial and vascular cells all critically interact to regulate the CBF fluctuations, both at rest and during functional hyperemia. In this regard, understanding how inflammation locally shapes cerebral hemodynamics is essential as this phenomenon is observed in many brain pathologies and drastically modifies the functional properties of the different components of the neurogliovascular unit (such as the induction of microglial and astrocytic reactivity). In addition to providing new elements for functional neuroimaging interpretation in patients, this may also lead to the identification of biomarkers or therapeutic perspectives.

Our main finding relates to the reorganization of cerebral hemodynamic activity towards an oscillatory state characterized by quasi-sinusoidal, high-amplitude oscillations at **∼**0.1 Hz, that locally propagate through the vascular tree as traveling waves under inflammation, as early as 24 h post-injection of LPS. This oscillatory mode is transient and is no longer detectable after seven days. Other studies have reported such sinusoidal hemodynamic oscillations at similar frequencies occurring sporadically in animals or humans under anesthesia^48,49^ or in the awake state^50^, but this has not yet been reported under the controlled induction of inflammation. The observed frequency is typical of two distinct phenomena, namely vasomotion and “Mayer waves”. The latter is due to regular fluctuations in blood pressure that can occur at around 0.1 Hz, which could obviously affect CBV changes^51^. However, as noted by other authors, such waves should be observed in all vessels simultaneously, whereas we and others have found these oscillations to be localized in discrete regions of the vasculature, propagating through the cortex - and in our case, occasionally between the cortex and subcortical regions - at a speed between 0.1 and 0.8 mm/s (previous reports^49,50^ and our own findings). Therefore, the most plausible phenomenon is vasomotion^52^, i.e. an autonomous and regular change in vessel diameter at about 0.1 Hz, of myogenic origin (through intracellular calcium fluctuations in smooth muscle cells^53^), distinct from the pulsatility due to the heartbeat. This is compatible with traveling waves, as vasomotion can propagate along the vessel walls thanks to gap-junctions and shear stress mechanisms^54^. A novel finding here is that vasomotor waves may also occur in subcortical areas and can travel long distances, for instance from the ventral hippocampus to the cortex in the contralateral hemisphere, which can be easily detected using fUSi as compared to previous studies describing hemodynamic oscillations using optical techniques. Recently, Broggini et al. also reported long-wavelength traveling waves in the cortical arterioles of awake mice due to vasomotion. Our findings align with their observation that such waves can span the entire cortical mantle^24^. Additionally, we reveal that the waves along the horizontal plane can propagate not only along the pial vessels, but also through adjacent penetrating cortical vessels. This results in hemodynamic changes that can span the entire thickness of cortex while traveling through distinct areas, not just the superficial layers. Importantly, we show here that fUSi is sensitive enough to track individual traveling waves, which calls for further mesoscopic studies of vasomotion in physiological or pathological conditions with this imaging modality.

Although vasomotion can occur in peripheral as well as cerebral vessels, it has previously been shown that cerebral vasomotion at 0.1 Hz is further entrained by changes in neuronal activity in the gamma-band, and this vaso-neuronal coupling would be the main driver of resting-state connectivity^7^. However, this has remained controversial as other works reported a lower correlation between vasomotion and electrophysiology measurements^8,48^. Here, we do confirm under pathological conditions that vasomotion can enhance homotopic functional connectivity, but whether these inflammatory hemodynamic oscillations are related or not to neuronal activity changes remains to be investigated. In this regard, we acknowledge that the absence of simultaneous electrophysiology measurements is an important limitation of our study that should be addressed in future investigations of the phenomenon. As functional hyperemia was not reduced in the presence of oscillations, we hypothesize that local neuronal activity was not strongly disrupted to the point of generating such sinusoidal hemodynamic waves by itself, as one might expect that a high basal neuronal activity in the absence of stimulation should result in a smaller difference of activity during the stimulation. Nevertheless, it is puzzling that the amplitude of functional hyperemia tended to be higher in the presence of strong vasomotion, and it suggests that the two phenomena can co-exist while engaging some common cellular and molecular mechanisms. The most plausible hypothesis to us is that vasomotion frequency is indeed controlled by internal vascular mechanisms and external neuronal activity, and vasomotion amplitude is further influenced by other players which are also involved in neurovascular coupling under functional hyperemia. Our data suggest that the missing piece of this puzzle is glial cells, particularly when they become reactive. In the same way, another study showed that non-reactive astrocytes are able to limit the amplitude of vasomotion by releasing prostaglandins during the constriction phase, which stimulate TRPV4 channels, increase Ca^2+^ in the terminal foot and activate COX-1^17^. As LPS has been shown to decrease the expression of COX-1 in reactive astrocytes^55^, this could increase the amplitude of vasomotion in our study by limiting negative feedback exerted by prostaglandins. In addition, there may be other mechanisms that reactive astrocytes could use to directly enhance vasomotion rather than reduce it. Moreover, while our findings highlight the role of glial reactivity, they again do not exclude a potential contribution from neurons. Astrocyte reactivity could secondarily impact neuronal homeostasis and excitability through disruptions in lactate metabolism and glutamate buffering^19^. In such a scenario, gliotoxin treatment would be expected to worsen oscillations rather than prevent them, due to stronger metabolic disruption and DAMP release from astrocyte injury^19,56^. Instead, chemically removing reactive astrocytes efficiently suppressed the amplification of hemodynamic oscillations, reinforcing the idea that reactive astrocytes—rather than general inflammation— are direct mediators of this phenomenon. Lastly, our findings also suggest that microglia might be either just necessary to locally initiate the astrocytic reactivity after being activated by LPS through TLR4 signaling, or directly involved in the amplification of hemodynamic oscillations, although this remains speculative and the possible molecular mechanisms are currently unclear.

Other interesting points should be discussed in terms of cellular changes in our model. Transcriptomics revealed a higher astrocytic reactivity at 48 h than at 7 days, with the identification of two astrocytic subclusters corresponding to a more basal state, characterized by moderate expression of *Gfap* and very low expression of *Nes*, including astrocytes from the VEH group and the LPS group at 7 days post-injection, and a strongly reactive state found only in the 48 h post-LPS group, with high expression of *Gfap* and *Nes*. This unbiased approach is recommended to take into account the complexity and heterogeneity of glial cells reactivity among different pathologies, as opposed to measuring a single marker^57–59^. Interestingly, nestin and GFAP are both intermediate filament proteins that are consistent with the known hypertrophy and morphological alterations occurring in reactive astrocytes, but *Nestin* increase has previously been reported in stroke and not following a peripheral injection of LPS in mice^60^, which indicates that the effects of LPS can vary significantly depending on the administration method. *Nestin* has also been described as absent from normal astrocytes and transiently expressed in reactive state, with a return to baseline levels at 7 days post- stroke^60^, which is similar to our own findings with LPS intracerebral injection in the rat cortex. Future works with optimized preparation protocols for enrichment of gliovascular unit should enable to better dissect the link between hemodynamic oscillations and astrocytic reactivity. Nevertheless, it is important to note that the intracerebral procedure itself could contribute to the differences observed between intracerebral and intraperitoneal LPS administration. The injury caused by needle insertion into the cortex may amplify the local effects of LPS, even though an intracerebral saline injection was used for the control group. This limitation of our model could also explain why the GFAP increase following LPS injection was not statistically significant compared to the control group, which tended to show already relatively high GFAP levels. Knowing the differences in half-life between *Gfap* mRNA (likely in the range of hours^61^) and GFAP protein (∼28 days^62^), the pro-inflammatory effects of the local injury may have transiently increased GFAP expression in the control group. While this effect may have already resolved at the mRNA level by 48 hours, leading to baseline *Gfap* levels in controls, the longer half-life of GFAP protein could have maintained elevated levels, leading to attenuated differences with the LPS group. An additional observation is that the morphology of microglial cells was slightly less impacted in the groups that received a co-injection of LPS and astrocytic toxins, suggesting that although microglia are the initial target of LPS effect through activation of the TLR4 pathway (with subsequent release of cytokines that render astrocytes reactive), reactive astrocytes may subsequently further increase the reactivity of microglia through a bidirectional interaction during inflammation^63,64^. This also illustrates the difficulty to isolate the specific contributions of microglia *versus* astrocytes under neuroinflammatory conditions. Finally, vascular cells (endothelial cells, smooth muscle vascular cells or pericytes) could also participate to increased vasomotion, either in direct interaction with reactive astrocytes or secondarily, through independent and additional pathways, given they are also dysregulated during inflammation^65^. Ongoing improvements in experimental protocols for specific manipulation of vascular cells^66^, as well as efficient isolation of these cells for transcriptomics^67^, will be important to investigate this possibility.

Regarding the effects of astrocytic toxins alone, we found that both tended to acutely increase functional hyperemia in the rat, with different effects that could be related to their different mechanisms of action. Surprisingly, AAA did not change the response in the visual cortex where it was injected, but the amplitude of response in the superior colliculus was significantly increased. The reason for this is unclear, although it might be possible that local acute perturbations of astrocytic activity in the visual cortex, and secondary neuronal activity, could alter inputs from the visual cortex to the superior colliculus^68^. The effect of FC was much more pronounced and resulted in a change of the shape of the response, with a sustained increase in CBV rather than a plateau at the end of the stimulation period. This is in line with a previous study showing that astrocytes are particularly important for the late component of functional hyperemia, observed during sustained sensory stimulation, such as in our own conditions^69^. Indeed, chemogenetic activation of astrocytes enhances late hyperemia in awake mice. We also found that FC induced such an effect, although we would expect it to reduce the response, since the toxin is supposed to reduce the calcium concentration in astrocytes and inhibit their activity^70^. This difference may be due to the use of anesthesia in our visual stimulation experiments, or to an acute compensatory hyperexcitability of astrocytes *in vivo* as opposed to the inhibitory effect of FC *in vitro*.

In terms of implications for functional imaging, the present study sheds new light on the complexity of cerebral hemodynamics and the involvement of glial cells in the regulation of cerebral blood flow. Such findings are important for understanding and interpreting the data, particularly in studies of patients with brain diseases. While future work will be needed to firmly demonstrate the causative influence of reactive glial cells, the present results suggest that vasomotion may serve as a hemodynamic biomarker for glial cell reactivity, enabling quantitative comparisons between patients/animal disease models using non-invasive imaging techniques through the measurement of frequency fluctuations between 0.08-0.15Hz, and with a high precision for mapping the territories displaying most glial reactivity as suggested by the spatial correlation results. This important perspective requires further validation by varying the injection site and comparing unilateral *versus* bilateral injections or evaluating other acute or chronic neuroinflammation models, as the functional properties of reactive glia are likely to vary significantly across brain regions and pathologies^57–59^. Notably, the sporadic reports of sinusoidal cerebral vasomotion in patients with brain tumors or during interictal periods in epilepsy (without insights into the underlying cellular mechanisms) demonstrates the clinical relevance of the phenomenon^49,50^. In practice, these results show that vasomotion and consequently hemodynamic waves can represent a huge part of the data variance (up to 99% in the LPS group). Whether frequencies above 0.08 Hz should be filtered out in fMRI resting-state analysis has been vastly debated^8,71^; here we argue that biologically relevant phenomena are occurring at such frequencies, providing insights into the glial components as well as our basic understanding of functional connectivity and hyperemia. Therefore, we suggest implementing separate analyzes of this frequency band, by simple measurements of the amplitude of frequency fluctuations, and by quantifying the traveling waves using peak detection, similarly to this paper (which could be further optimized as our method was quite stringent regarding the waves classification), or complex PCA^72^ to increase the interpretability of future functional neuroimaging studies.

Finally, although vasomotion is still a poorly understood phenomenon, it has been suggested to facilitate oxygen exchange between vessels and the tissues^52^. As hypoxia is frequently observed in inflammatory conditions^73^ and LPS can activate the transcription factor hypoxia-inducible factor 1 (HIF-1)^74^, the present effect could compensate neuronal hypoxia in pathological conditions. Importantly, it has been shown that an “artificial” increase in cerebral vasomotion by functional hyperemia improves the clearance of waste products along the vessels^75^ and several studies have emphasized that this should be a major driving force for improving exchanges between the cerebrospinal (CSF) and interstitial (ISF) fluids, according to the glymphatic system hypothesis^76^ as well as the intra-mural peri-arterial drainage hypothesis^77,78^. Although this remains speculative at this early stage, it is conceivable that in inflammatory conditions, enhancement of vasomotion may improve CSF/ISF exchange to enhance waste removal, or even facilitate the communication between the CSF/ISF and the meningeal lymphatic system^79^. This may explain the reported paradoxical improvement in soluble amyloid beta-peptide following intracerebral injection of LPS in the mice^80^. These different possibilities suggest new therapeutic opportunities by modulating the cerebral vasomotion in a pathological context such as neurodegenerative disorders, and deserve further investigation.

## Contributors

MP, MD, LZ and BV conceived the project. MP, MV and BV performed the imaging experiments and surgeries. MP, LV and BV analyzed the fUSi and ULM data. MP performed the immunofluorescence staining and analysis. GM performed the single-nuclei RNA-seq experiment and analysis. SB, PM, JCC and OP designed and performed the analysis of microglia morphology. MD, LZ and BV supervised the project. MP and BV wrote the first draft. MP, MD, LZ and BV reviewed the article. MP and BV have accessed and verified the underlying data. All authors read and approved the final manuscript.

## Acknowledgments

This work was funded by the ANR (LabCom “Neuroimaging for Drug Discovery”) and the Auvergne-Rhône-Alpes Region (Pack Ambition Recherche “Brain Imaging for Drug Discovery”). We thank the CIQLE facility and Bruno Chapuis for the assistance with the slide scanner, the Cancer Genomic Platform (PGC) and Cyril Dégletagne at the Centre de Recherche en Cancérologie de Lyon (CRCL) for help in producing the transcriptional dataset, and Jean-Charles Mariani and Samuel Le Meur-Diebolt for insightful discussions and help with BIDS standardization of the fUSi dataset.

## Data sharing statement

Unprocessed fUSi images (following the BIDS standardization extension proposal for fUSi) are available at Zenodo (doi: 10.5281/zenodo.15194839). The single-nuclei transcriptomic data are available in the GEO database under the accession number GSE292745. Other data will be made available upon request to the authors.

## REFERENCES

1. Howarth C, Mishra A, Hall CN. More than just summed neuronal activity: how multiple cell types shape the BOLD response. Phil Trans R Soc B. 2021 Jan 4;376(1815):20190630.

2. Mishra A, Hall CN, Howarth C, Freeman RD. Key relationships between non-invasive functional neuroimaging and the underlying neuronal activity. Phil Trans R Soc B. 2021 Jan 4;376(1815):20190622.

3. Villringer A. Understanding Functional Neuroimaging Methods Based on Neurovascular Coupling. In: Villringer A, Dirnagl U, editors. Optical Imaging of Brain Function and Metabolism 2. Boston, MA: Springer US; 1997. p. 177–93. (Advances in Experimental Medicine and Biology; vol. 413).

4. Masamoto K, Vazquez A. Optical imaging and modulation of neurovascular responses. J Cereb Blood Flow Metab. 2018 Dec;38(12):2057–72.

5. Liao LD, Tsytsarev V, Delgado-Martínez I, Li ML, Erzurumlu R, Vipin A, et al. Neurovascular coupling: in vivo optical techniques for functional brain imaging. BioMed Eng OnLine. 2013;12(1):38.

6. Deffieux T, Demené C, Tanter M. Functional Ultrasound Imaging: A New Imaging Modality for Neuroscience. Neuroscience. 2021 Oct;474:110–21.

7. Mateo C, Knutsen PM, Tsai PS, Shih AY, Kleinfeld D. Entrainment of Arteriole Vasomotor Fluctuations by Neural Activity Is a Basis of Blood-Oxygenation-Level-Dependent “Resting-State” Connectivity. Neuron. 2017 Nov;96(4):936–948.e3.

8. Lambers H, Wachsmuth L, Lippe C, Faber C. The impact of vasomotion on analysis of rodent fMRI data. Front Neurosci. 2023 Feb 24;17:1064000.

9. Mishra A, Gordon GR, MacVicar BA, Newman EA. Astrocyte Regulation of Cerebral Blood Flow in Health and Disease. Cold Spring Harb Perspect Biol. 2024 Feb 5;a041354.

10. Nippert AR, Biesecker KR, Newman EA. Mechanisms Mediating Functional Hyperemia in the Brain. Neuroscientist. 2018 Feb;24(1):73–83.

11. Howarth C. The contribution of astrocytes to the regulation of cerebral blood flow. Front Neurosci. 2014 May 9;8.

12. Petzold GC, Murthy VN. Role of Astrocytes in Neurovascular Coupling. Neuron. 2011 Sep;71(5):782–97.

13. Bazargani N, Attwell D. Astrocyte calcium signaling: the third wave. Nat Neurosci. 2016 Feb;19(2):182–9.

14. Lia A, Di Spiezio A, Speggiorin M, Zonta M. Two decades of astrocytes in neurovascular coupling. Front Netw Physiol. 2023 Apr 3;3:1162757.

15. MacVicar BA, Newman EA. Astrocyte Regulation of Blood Flow in the Brain. Cold Spring Harb Perspect Biol. 2015 May;7(5):a020388.

16. Filosa JA, Bonev AD, Straub SV, Meredith AL, Wilkerson MK, Aldrich RW, et al. Local potassium signaling couples neuronal activity to vasodilation in the brain. Nat Neurosci. 2006 Nov;9(11):1397–403.

17. Haidey JN, Peringod G, Institoris A, Gorzo KA, Nicola W, Vandal M, et al. Astrocytes regulate ultra-slow arteriole oscillations via stretch-mediated TRPV4-COX-1 feedback. Cell Reports. 2021 Aug;36(5):109405.

18. Császár E, Lénárt N, Cserép C, Környei Z, Fekete R, Pósfai B, et al. Microglia modulate blood flow, neurovascular coupling, and hypoperfusion via purinergic actions. Journal of Experimental Medicine. 2022 Mar 7;219(3):e20211071.

19. Verkhratsky A, Butt A, Li B, Illes P, Zorec R, Semyanov A, et al. Astrocytes in human central nervous system diseases: a frontier for new therapies. Sig Transduct Target Ther. 2023 Oct 13;8(1):396.

20. Boido D, Rungta RL, Osmanski BF, Roche M, Tsurugizawa T, Le Bihan D, et al. Mesoscopic and microscopic imaging of sensory responses in the same animal. Nat Commun. 2019 Mar 7;10(1):1110.

21. Urban A, Dussaux C, Martel G, Brunner C, Mace E, Montaldo G. Real-time imaging of brain activity in freely moving rats using functional ultrasound. Nat Methods. 2015 Sep;12(9):873–8.

22. El Hady A, Takahashi D, Sun R, Akinwale O, Boyd-Meredith T, Zhang Y, et al. Chronic brain functional ultrasound imaging in freely moving rodents performing cognitive tasks. Journal of Neuroscience Methods. 2024 Mar;403:110033.

23. Tiran E, Ferrier J, Deffieux T, Gennisson JL, Pezet S, Lenkei Z, et al. Transcranial Functional Ultrasound Imaging in Freely Moving Awake Mice and Anesthetized Young Rats without Contrast Agent. Ultrasound in Medicine & Biology. 2017 Aug;43(8):1679–89.

24. Broggini T, Duckworth J, Ji X, Liu R, Xia X, Mächler P, et al. Long-wavelength traveling waves of vasomotion modulate the perfusion of cortex. Neuron. 2024 Jul;112(14):2349–2367.e8.

25. Osmanski BF, Pezet S, Ricobaraza A, Lenkei Z, Tanter M. Functional ultrasound imaging of intrinsic connectivity in the living rat brain with high spatiotemporal resolution. Nat Commun. 2014 Oct 3;5(1):5023.

26. Urban A, Mace E, Brunner C, Heidmann M, Rossier J, Montaldo G. Chronic assessment of cerebral hemodynamics during rat forepaw electrical stimulation using functional ultrasound imaging. NeuroImage. 2014 Nov;101:138–49.

27. Bercoff J, Montaldo G, Loupas T, Savery D, Mézière F, Fink M, et al. Ultrafast compound doppler imaging: providing full blood flow characterization. IEEE Trans Ultrason, Ferroelect, Freq Contr. 2011 Jan;58(1):134–47.

28. Montaldo G, Tanter M, Bercoff J, Benech N, Fink M. Coherent plane-wave compounding for very high frame rate ultrasonography and transient elastography. IEEE Trans Ultrason, Ferroelect, Freq Contr. 2009 Mar;56(3):489–506.

29. Demene C, Deffieux T, Pernot M, Osmanski BF, Biran V, Gennisson JL, et al. Spatiotemporal Clutter Filtering of Ultrafast Ultrasound Data Highly Increases Doppler and fUltrasound Sensitivity. IEEE Trans Med Imaging. 2015 Nov;34(11):2271–85.

30. Grandjean J, Desrosiers-Gregoire G, Anckaerts C, Angeles-Valdez D, Ayad F, Barrière DA, et al. A consensus protocol for functional connectivity analysis in the rat brain. Nat Neurosci. 2023 Apr;26(4):673–81.

31. Droguerre M, Vidal B, Valdebenito M, Mouthon F, Zimmer L, Charvériat M. Impaired Local and Long-Range Brain Connectivity and Visual Response in a Genetic Rat Model of Hyperactivity Revealed by Functional Ultrasound. Front Neurosci. 2022 Mar 24;16:865140.

32. Gesnik M, Blaize K, Deffieux T, Gennisson JL, Sahel JA, Fink M, et al. 3D functional ultrasound imaging of the cerebral visual system in rodents. NeuroImage. 2017 Apr;149:267–74.

33. Errico C, Pierre J, Pezet S, Desailly Y, Lenkei Z, Couture O, et al. Ultrafast ultrasound localization microscopy for deep super-resolution vascular imaging. Nature. 2015 Nov;527(7579):499–502.

34. Vidal B, Pereira M, Valdebenito M, Vidal L, Mouthon F, Zimmer L, et al. Pharmaco-fUS in cognitive impairment: Lessons from a preclinical model. J Psychopharmacol. 2022 Nov;36(11):1273–9.

35. Vidal B, Droguerre M, Venet L, Valdebenito M, Mouthon F, Zimmer L, et al. Inter-subject registration and application of the SIGMA rat brain atlas for regional labeling in functional ultrasound imaging. Journal of Neuroscience Methods. 2021 May;355:109139.

36. Barrière DA, Magalhães R, Novais A, Marques P, Selingue E, Geffroy F, et al. The SIGMA rat brain templates and atlases for multimodal MRI data analysis and visualization. Nat Commun. 2019 Dec 13;10(1):5699.

37. Song XW, Dong ZY, Long XY, Li SF, Zuo XN, Zhu CZ, et al. REST: A Toolkit for Resting-State Functional Magnetic Resonance Imaging Data Processing. Harrison BJ, editor. PLoS ONE. 2011 Sep 20;6(9):e25031.

38. Cabral J, Fernandes FF, Shemesh N. Intrinsic macroscale oscillatory modes driving long range functional connectivity in female rat brains detected by ultrafast fMRI. Nat Commun. 2023 Feb 6;14(1):375.

39. Wang Y, DelRosso NV, Vaidyanathan TV, Cahill MK, Reitman ME, Pittolo S, et al. Accurate quantification of astrocyte and neurotransmitter fluorescence dynamics for single-cell and population-level physiology. Nat Neurosci. 2019 Nov;22(11):1936–44.

40. Benkeder S, Dinh SM, Marchal P, Gea PD, Thoby-Brisson M, Hubert V, et al. MorphoCellSorter: An Andrews plot-based sorting approach to rank microglia according to their morphological features. eLife. 2024. Available from: https://elifesciences.org/reviewed-preprints/101630v1

41. Yu G, Wang LG, Han Y, He QY. clusterProfiler: an R Package for Comparing Biological Themes Among Gene Clusters. OMICS: A Journal of Integrative Biology. 2012 May;16(5):284–7.

42. Khurgel M, Koo AC, Ivy GO. Selective ablation of astrocytes by intracerebral injections ofD α-aminoadipate. Glia. 1996 Apr;16(4):351–8.

43. F Fonnum F, Johnsen A, Hassel B. Use of fluorocitrate and fluoroacetate in the study of brain metabolism. Glia. 1997 Sep;21(1):106–13.

44. Hassel B, Paulsen RE, Johnsen A, Fonnum F. Selective inhibition of glial cell metabolism in vivo by fluorocitrate. Brain Research. 1992 Mar;576(1):120–4.

45. Zhang Z, Ma Z, Zou W, Guo H, Liu M, Ma Y, et al. The Appropriate Marker for Astrocytes: Comparing the Distribution and Expression of Three Astrocytic Markers in Different Mouse Cerebral Regions. BioMed Research International. 2019 Jun 24;2019:1–15.

46. Boche D, Perry VH, Nicoll JAR. Review: Activation patterns of microglia and their identification in the human brain. Neuropathology Appl Neurobio. 2013 Feb;39(1):3–18.

47. Figley CR, Stroman PW. The role(s) of astrocytes and astrocyte activity in neurometabolism, neurovascular coupling, and the production of functional neuroimaging signals: Astrocyte control of fMRI and PET signals. European Journal of Neuroscience. 2011 Feb;33(4):577–88.

48. Rivadulla C, De Labra C, Grieve KL, Cudeiro J. Vasomotion and Neurovascular Coupling in the Visual Thalamus In Vivo. Dickson CT, editor. PLoS ONE. 2011 Dec 9;6(12):e28746.

49. Noordmans HJ, Van Blooijs D, Siero JCW, Zwanenburg JJM, Klaessens JHGM, Ramsey NF. Detailed view on slow sinusoidal, hemodynamic oscillations on the human brain cortex by Fourier transforming oxy/deoxy hyperspectral images. Human Brain Mapping. 2018 Sep;39(9):3558–73.

50. Rayshubskiy A, Wojtasiewicz TJ, Mikell CB, Bouchard MB, Timerman D, Youngerman BE, et al. Direct, intraoperative observation of ∼0.1 Hz hemodynamic oscillations in awake human cortex: implications for fMRI. Neuroimage. 2014 Feb 15;87:323–31.

51. Julien C. The enigma of Mayer waves: Facts and models. Cardiovascular Research. 2006 Apr 1;70(1):12–21.

52. Aalkjær C, Boedtkjer D, Matchkov V. Vasomotion – what is currently thought? Acta Physiologica. 2011 Jul;202(3):253–69.

53. Li J, Zhang Y, Zhang D, Wang W, Xie H, Ruan J, et al. Ca2+ oscillation in vascular smooth muscle cells control myogenic spontaneous vasomotion and counteract post-ischemic no-reflow. Commun Biol. 2024 Mar 15;7(1):332.

54. Koenigsberger M, Sauser R, Bény JL, Meister JJ. Effects of Arterial Wall Stress on Vasomotion. Biophysical Journal. 2006 Sep;91(5):1663–74.

55. Font-Nieves M, Sans-Fons MG, Gorina R, Bonfill-Teixidor E, Salas-Pérdomo A, Márquez-Kisinousky L, et al. Induction of COX-2 Enzyme and Down-regulation of COX-1 Expression by Lipopolysaccharide (LPS) Control Prostaglandin E2 Production in Astrocytes. Journal of Biological Chemistry. 2012 Feb;287(9):6454–68.

56. Rosciszewski G, Cadena V, Murta V, Lukin J, Villarreal A, Roger T, et al. Toll-Like Receptor 4 (TLR4) and Triggering Receptor Expressed on Myeloid Cells-2 (TREM-2) Activation Balance Astrocyte Polarization into a Proinflammatory Phenotype. Mol Neurobiol [Internet]. 2017 May 25; Available from: http://link.springer.com/10.1007/s12035-017-0618-z

57. Hasel P, Aisenberg WH, Bennett FC, Liddelow SA. Molecular and metabolic heterogeneity of astrocytes and microglia. Cell Metabolism. 2023 Apr;35(4):555–70.

58. Sun J, Song Y, Chen Z, Qiu J, Zhu S, Wu L, et al. Heterogeneity and Molecular Markers for CNS Glial Cells Revealed by Single-Cell Transcriptomics. Cell Mol Neurobiol. 2022 Nov;42(8):2629–42.

59. Escartin C, Galea E, Lakatos A, O’Callaghan JP, Petzold GC, Serrano-Pozo A, et al. Reactive astrocyte nomenclature, definitions, and future directions. Nat Neurosci. 2021 Mar;24(3):312–25.

60. Zamanian JL, Xu L, Foo LC, Nouri N, Zhou L, Giffard RG, et al. Genomic Analysis of Reactive Astrogliosis. J Neurosci. 2012 May 2;32(18):6391–410.

61. Condorelli DF, Dell’Albani P, Kaczmarek L, Messina L, Spampinato G, Avola R, et al. Glial fibrillary acidic protein messenger RNA and glutamine synthetase activity after nervous system injury. J of Neuroscience Research. 1990 Jun;26(2):251–7.

62. Moody LR, Barrett-Wilt GA, Sussman MR, Messing A. Glial fibrillary acidic protein exhibits altered turnover kinetics in a mouse model of Alexander disease. Journal of Biological Chemistry. 2017 Apr;292(14):5814–24.

63. RuedaDCarrasco J, MartinDBermejo MJ, Pereyra G, Mateo MI, Borroto A, Brosseron F, et al. SFRP1 modulates astrocyteDtoDmicroglia crosstalk in acute and chronic neuroinflammation. EMBO Reports. 2021 Nov 4;22(11):e51696.

64. Pascual O, Ben Achour S, Rostaing P, Triller A, Bessis A. Microglia activation triggers astrocyte-mediated modulation of excitatory neurotransmission. Proc Natl Acad Sci USA. 2012 Jan 24;109(4).

65. Obadia N, Andrade G, Leardini-Tristão M, Albuquerque L, Garcia C, Lima F, et al. TLR4 mutation protects neurovascular function and cognitive decline in high-fat diet-fed mice. J Neuroinflammation. 2022 Apr 29;19(1):104.

66. Abe Y, Kwon S, Oishi M, Unekawa M, Takata N, Seki F, et al. Optical manipulation of local cerebral blood flow in the deep brain of freely moving mice. Cell Reports. 2021 Jul;36(4):109427.

67. Bjørnholm KD, Del Gaudio F, Li H, Li W, Vazquez-Liebanas E, Mäe MA, et al. A robust and efficient microvascular isolation method for multimodal characterization of the mouse brain vasculature. Cell Reports Methods. 2023 Mar;3(3):100431.

68. Zhao X, Liu M, Cang J. Visual Cortex Modulates the Magnitude but Not the Selectivity of Looming-Evoked Responses in the Superior Colliculus of Awake Mice. Neuron. 2014 Oct;84(1):202–13.

69. Institoris A, Vandal M, Peringod G, Catalano C, Tran CH, Yu X, et al. Astrocytes amplify neurovascular coupling to sustained activation of neocortex in awake mice. Nat Commun. 2022 Dec 22;13(1):7872.

70. Padmashri R, Suresh A, Boska MD, Dunaevsky A. Motor-Skill Learning Is Dependent on Astrocytic Activity. Neural Plasticity. 2015;2015:1–11.

71. Davey CE, Grayden DB, Egan GF, Johnston LA. Filtering induces correlation in fMRI resting state data. NeuroImage. 2013 Jan;64:728–40.

72. Bolt T, Nomi JS, Bzdok D, Salas JA, Chang C, Thomas Yeo BT, et al. A parsimonious description of global functional brain organization in three spatiotemporal patterns. Nat Neurosci. 2022 Aug;25(8):1093–103.

73. Kölliker-Frers R, Udovin L, Otero-Losada M, Kobiec T, Herrera MI, Palacios J, et al. Neuroinflammation: An Integrating Overview of Reactive-Neuroimmune Cell Interactions in Health and Disease. Giovarelli M, editor. Mediators of Inflammation. 2021 May 31;2021:1–20.

74. Frede S, Stockmann C, Freitag P, Fandrey J. Bacterial lipopolysaccharide induces HIF-1 activation in human monocytes via p44/42 MAPK and NF-κB. Biochemical Journal. 2006 Jun 15;396(3):517–27.

75. Van Veluw SJ, Hou SS, Calvo-Rodriguez M, Arbel-Ornath M, Snyder AC, Frosch MP, et al. Vasomotion as a Driving Force for Paravascular Clearance in the Awake Mouse Brain. Neuron. 2020 Feb;105(3):549–561.e5.

76. Hablitz LM, Nedergaard M. The Glymphatic System: A Novel Component of Fundamental Neurobiology. J Neurosci. 2021 Sep 15;41(37):7698–711.

77. Diem AK, Carare RO, Weller RO, Bressloff NW. A control mechanism for intra-mural peri-arterial drainage via astrocytes: How neuronal activity could improve waste clearance from the brain. Deli MA, editor. PLoS ONE. 2018 Oct 4;13(10):e0205276.

78. Aldea R, Weller RO, Wilcock DM, Carare RO, Richardson G. Cerebrovascular Smooth Muscle Cells as the Drivers of Intramural Periarterial Drainage of the Brain. Front Aging Neurosci. 2019 Jan 23;11:1.

79. Da Mesquita S, Fu Z, Kipnis J. The Meningeal Lymphatic System: A New Player in Neurophysiology. Neuron. 2018 Oct;100(2):375–88.

80. DiCarlo G. Intrahippocampal LPS injections reduce AÎ^2^ load in APP+PS1 transgenic mice. Neurobiology of Aging. 2001 Dec;22(6):1007–12.

