## Supplementary figures and images for "Induction of hemodynamic traveling waves by glial-related vasomotion in a rat model of neuroinflammation: implications for functional neuroimaging"

### Supplementary Figure S1

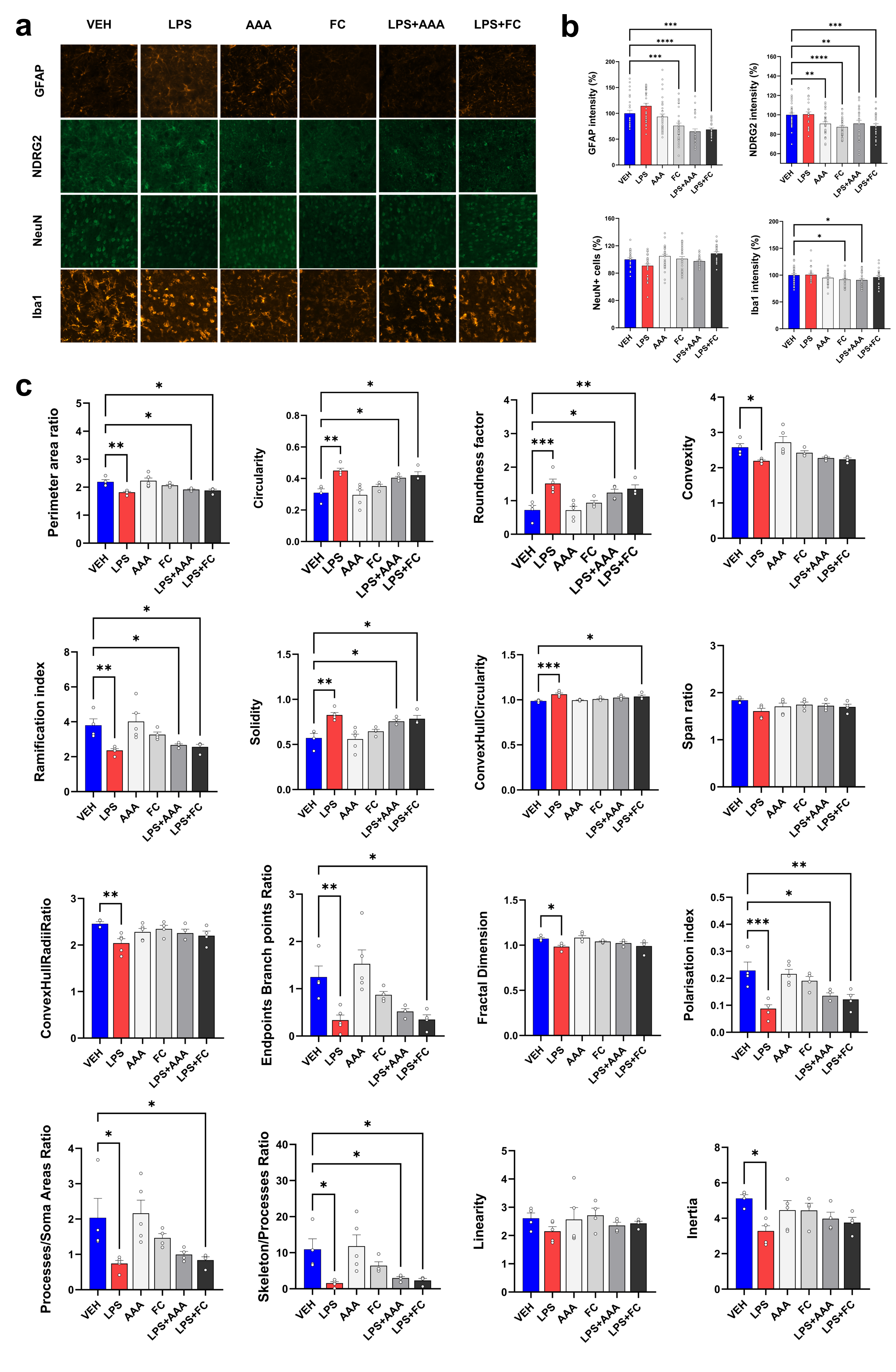

### Supplementary Figure S2

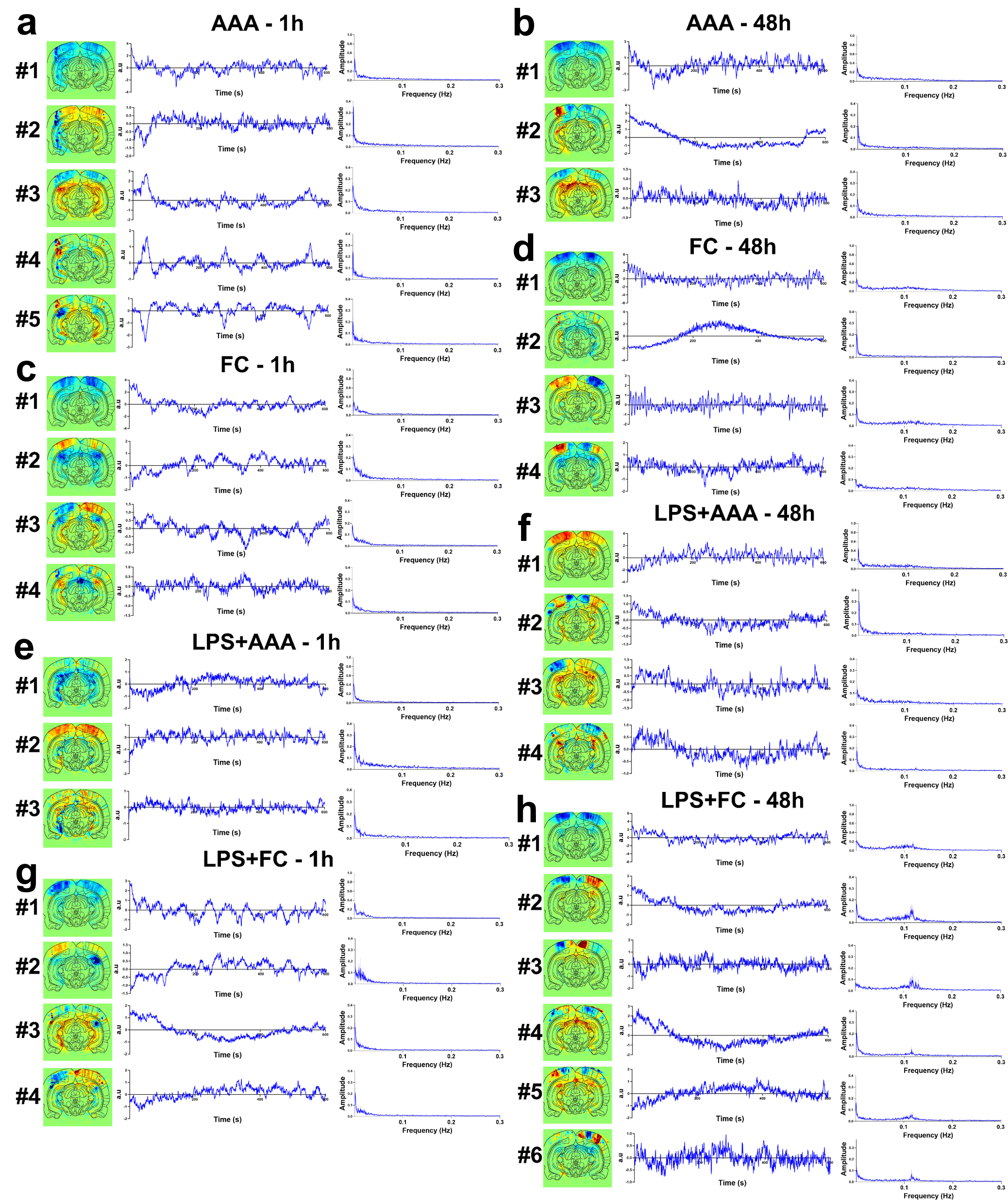

### Supplementary Figure S3

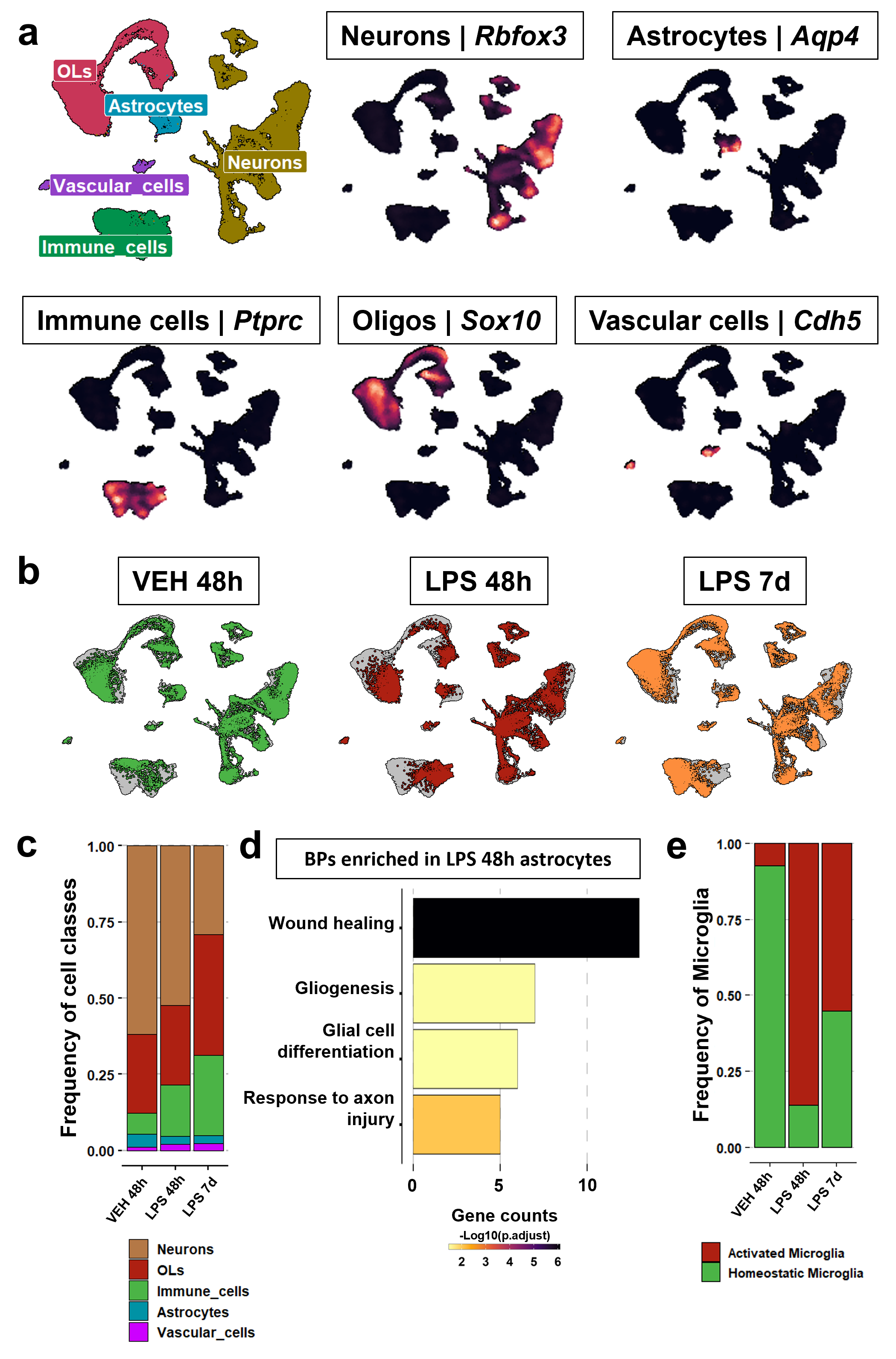

### Supplementary Figure S4

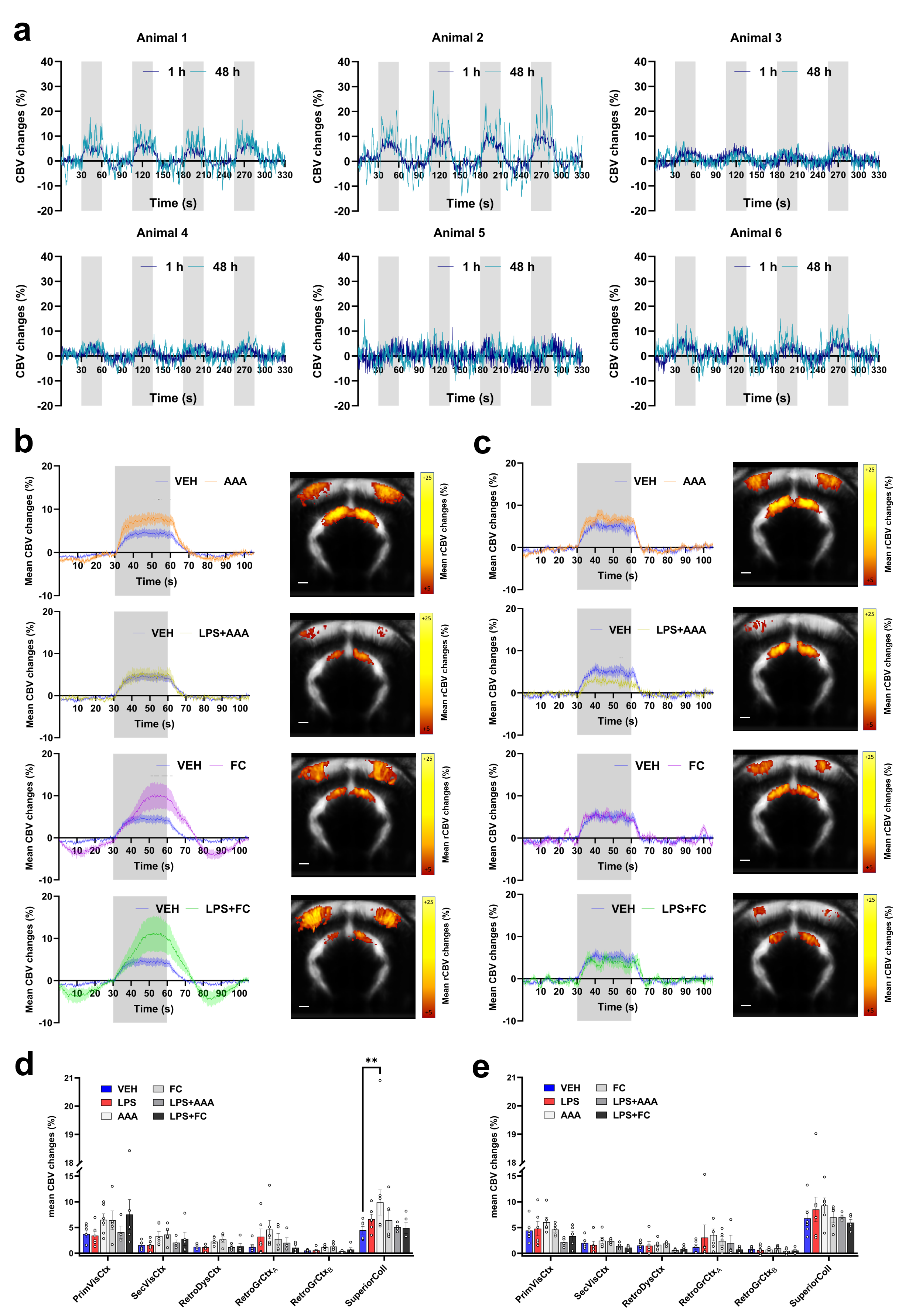

### Supplementary Figure S5

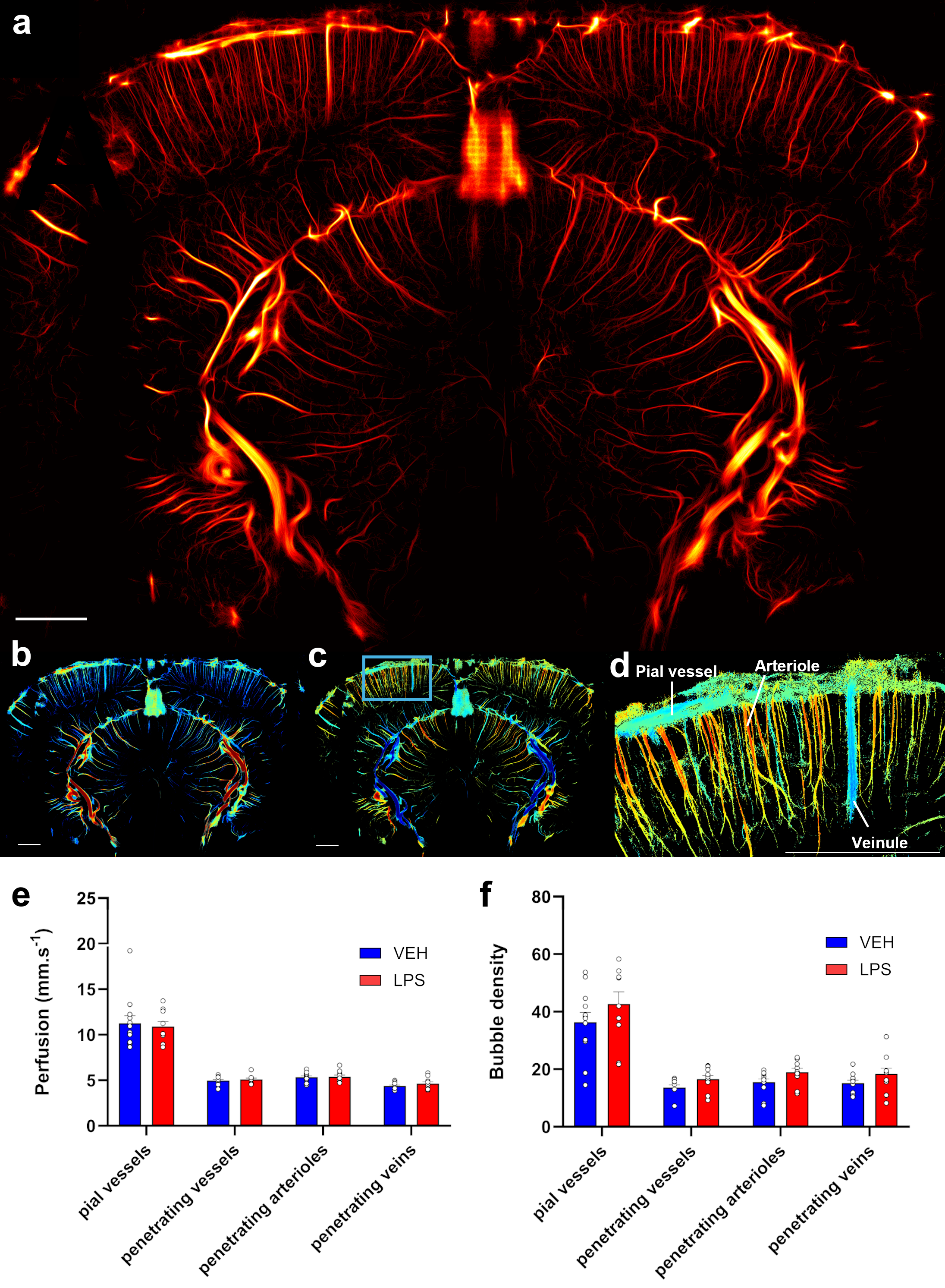
