## Supplementary Table 1 for "Induction of hemodynamic traveling waves by glial-related vasomotion in a rat model of neuroinflammation: implications for functional neuroimaging"

| **Marker** | **Supplier** | **RRID tag** | **Species** | **Dilution** |
| --- | --- | --- | --- | --- |
| **PRIMARY ANTIBODIES** | | | | |
| GFAP | Dako (Z0334) | AB_10013382 | Rabbit | 1:200 |
| NDRG2 | Santa Cruz (sc-376202) | AB_11008209 | Mouse | 1:500 |
| NeuN | Merck – (MAB377) | AB_2298772 | Mouse | 1:1000 |
| Iba1 | Synaptic Systems – (234 003) | AB_10641962 | Rabbit | 1:1000 |
| **SECONDARY ANTIBODIES** | | | | |
| Dylight 488  (Anti-rabbit) | Invitrogen – (35503) | AB_1965946 | Goat | 1:250 |
| Alexa Fluor 546  (Anti-mouse) | Thermo Fisher – (A11010) | AB_2534077 | Goat | 1:500 |
